# Single-cell profiling and SCOPE-seq reveal the lineage dynamics of adult neurogenesis and NOTUM as a key V-SVZ regulator

**DOI:** 10.1101/770610

**Authors:** Dogukan Mizrak, N. Sumru Bayin, Jinzhou Yuan, Zhouzerui Liu, Radu Suciu, Micah J. Niphakis, Nhi Ngo, Kenneth M. Lum, Benjamin F. Cravatt, Alexandra L. Joyner, Peter A. Sims

## Abstract

Neural stem cells (NSCs) and their progeny reside in specialized niches in the adult mammalian brain where they generate new neurons and glia throughout life. Adult NSCs of the ventricular-subventricular zone (V-SVZ) are prone to rapid exhaustion; thus timely, context-dependent neurogenesis demands adaptive signaling among the vast number of neighboring progenitors nestled between the ventricular surface and nearby blood vessels. To dissect adult neuronal lineage progression and regulation, we profiled >56,000 V-SVZ and olfactory bulb (OB) cells by single-cell RNA-sequencing (scRNA-seq). Our analyses revealed the diversity of V-SVZ-derived OB neurons, the temporal dynamics of lineage progression, and a key intermediate NSC population enriched for expression of *Notum*, which encodes a secreted WNT antagonist. Single Cell Optical Phenotyping and Expression (SCOPE-seq), a technology linking live cell imaging with scRNA-seq, uncovered dynamic control of cell size concomitant with NSC differentiation with *Notum*^+^ NSCs at a critical size poised for cell division, and a preference of NOTUM surface binding to neuronal precursors with active WNT signaling. Finally, *in vivo* pharmacological inhibition of NOTUM significantly expanded neuronal precursor pools in the V-SVZ. Our findings highlight a critical regulatory state during NSC activation marked by NOTUM, a secreted enzyme that ensures efficient neurogenesis by preventing WNT signaling activation in NSC progeny.

## INTRODUCTION

Neurogenesis persists in two principal regions of the adult mouse brain: the subgranular zone of the dentate gyrus and the V-SVZ located in the walls of the lateral ventricles (Doetsch et al., 1999; Fiorelli et al., 2015; Mirzadeh et al., 2008). The V-SVZ contains both quiescent (qNSCs) and actively dividing NSCs (aNSCs) (Codega et al., 2014; Mich et al., 2014) with intrinsic regional identities defined during embryogenesis (Fuentealba et al., 2015), generating different OB interneuron subtypes or glia depending on their location (Brill et al., 2009; Delgado and Lim, 2015; Kohwi et al., 2005; Lopez-Juarez et al., 2013; Merkle et al., 2014; Merkle et al., 2007; Mizrak et al., 2019; Ortega et al., 2013; Zweifel et al., 2018). V-SVZ qNSCs are specialized astrocytes with radial morphology (reviewed in Kriegstein and Alvarez-Buylla, 2009; Mirzadeh et al., 2008), and once activated divide symmetrically, resulting in their depletion over time (Obernier et al., 2018). aNSCs give rise to transit-amplifying cells (TACs), which generate neuroblasts (NBs). NBs migrate via the rostral migratory stream (RMS) to the OB (Figure 1A), where they terminally differentiate into interneurons. V-SVZ NSC-to-OB interneuron differentiation requires cells with a broad distribution of developmental states; however, the identities and the functions of key intermediate states and the gene expression programs pertinent to these transitions are unknown. While recent efforts have defined OB neuron subtypes (Eng et al., 2019; Tepe et al., 2018), classification of V-SVZ-derived OB neurons by fate-mapped scRNA-seq is still lacking.

**Figure 1.**
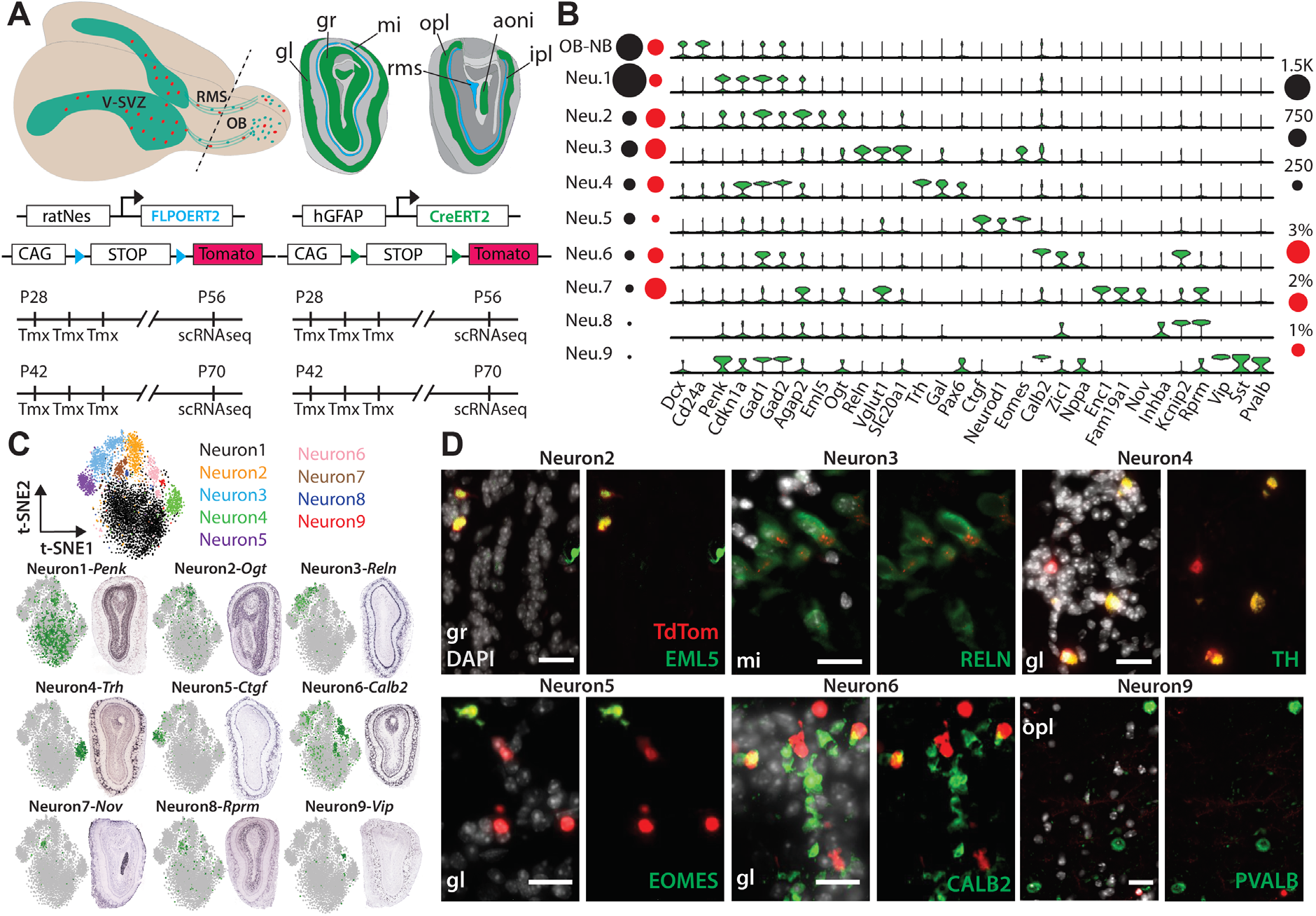
Fate-mapping experiments and OB Neuron Subtypes. **(A)** Top: Schema of adult mouse brain (left) showing the migration of V-SVZ neuronal precursors (red) to the OB. Coronal schema (right) highlighting the multi-layered architecture of the OB (gl: glomerular layer, gr: granule cell layer, mi: mitral cell layer, epl: external plexiform layer, ipl: internal plexiform layer, aoni: anterior olfactory nucleus, rms/sez: rostral migratory stream which terminates in the subependymal zone (SEZ) of OB). Bottom: Four and six-week old GCERT2 and NESFLPO animals were injected with tamoxifen for three consecutive days and assayed four weeks after the last injection. **(B)** Violin plots showing log_2_(CPM+1) (counts per million) expression values of neuron subtype markers (FDR<0.05). Black and red circles indicate cell numbers and fraction of reporter-positive cells, respectively, in each cluster. **(C)** Top: t-distributed stochastic neighbor embedding (t-SNE) projections of major OB cell types colored separately. Bottom: ISH images showing spatial distribution of subtype markers and their scaled expression on t-SNE. **(D)** Coronal images showing endogenous TdTom and main OB neuron subtype marker co-localization. The corresponding OB layer was indicated on each image. TdTom expression in *mi* layer RELN+ cells was noticeably weaker. Scale bars: 20μm.

Intercellular communication in the V-SVZ is essential for proper adult neurogenesis, but has not been fully explored, particularly at the single-cell level. GABA released by NBs inhibits NSC proliferation (Alfonso et al., 2012; Liu et al., 2005), while contact mediated NOTCH signaling between NSCs and TACs regulates NSC quiescence (Kawaguchi et al., 2013). WNT signaling is also known to play a crucial role in orchestrating proliferation and differentiation of adult V-SVZ NSCs (Adachi et al., 2007; Azim et al., 2014; Ortega et al., 2013; Qu et al., 2010; Zywitza et al., 2018). Although these studies provide valuable insights into the significance of WNT signaling in V-SVZ, they do not reveal the key WNT signaling regulators, or a mechanistic understanding of intercellular WNT signaling interactions. Importantly, WNT signaling can be negatively regulated by multiple families of secreted proteins that modulate WNT ligand affinity for the Frizzled receptors of the WNT signaling pathway (reviewed in Driehuis and Clevers, 2017). Therefore; these secreted negative regulators could facilitate adaptive signaling between neuronal progenitors to cope with sustained WNT ligand exposure.

Highly scalable scRNA-seq technologies (Klein et al., 2015; Macosko et al., 2015; Yuan and Sims, 2016; Zheng et al., 2016) have been applied to address cellular heterogeneity and temporal gene expression changes in various stem cell contexts including the V-SVZ field (Mizrak et al., 2019; Tepe et al., 2018; Zywitza et al., 2018); however these V-SVZ studies lacked genetically-induced fate-mapping reporters and could not link V-SVZ NSCs to their progeny in the olfactory bulb. Importantly, current high-throughput methods are unable to link live cell imaging and scRNA-seq data from the same cell, and cannot measure, for example, morphological features of individual cells. Moreover, due to the low sensitivity of large-scale scRNA-seq technologies, many of the expressed transcripts in a given cell are undetected. This is particularly problematic in applications of these technologies to stem cells, where an intracellular reporter or a surface marker is often used to label cells. In this study, we applied large-scale scRNA-seq and an improved version of SCOPE-seq (Yuan et al., 2018), which directly links live-cell imaging and scRNA-seq, to reveal the cellular dynamics of adult neurogenesis. Importantly, we uncovered a rare intermediate population marked by NOTUM, which we demonstrate to be a key mediator of V-SVZ intercellular signaling.

## RESULTS

### Large-Scale Single Cell Profiling of Adult Mouse Olfactory Bulb with Fate-mapping Reporters

We profiled >56,000 cells from matched V-SVZs and OBs of animals carrying either the well-established *hGFAP::CreERT2; Rosa26^LSL-TdTomato^* (GCERT2) (Ganat et al., 2006; Mich et al., 2014; Obernier et al., 2018) or a newer V-SVZ enriched *ratNes::FLPOER; Rosa26^FSF-TdTomato^* (NESFLPO) fate-mapping reporter (Lao et al., 2012; Wojcinski et al., 2017) (**Figure 1A, Figure S1A**). NESFLPO specifically labels cells on lateral ventricle walls, and robustly generates *TdTomato*^+^ (*TdTom*^+^) OB cells four weeks after tamoxifen induction (**Figure S1B**). We identified ten major OB cell populations including migrating NBs and mature neurons (Levine et al., 2015; Shekhar et al., 2016) (**Figure S1C**). Subclustering of the >4,500 OB neurons revealed nine molecularly distinct subtypes (**Figure 1B, Table S1**), although morphological classification of OB neurons is more complex and dependent on overlapping markers (reviewed in Nagayama et al., 2014). *TdTom* expression was detected in multiple neuron subtypes (64 *TdTom*^+^ neurons), with ≥50% of *TdTom*^+^ neurons in the abundant Neuron1 and 2 (34 *TdTom*^+^ neurons), while the fraction of *TdTom*^+^ neurons was significantly enriched in Neuron3 and 7 (**Figure 1B**). *Gad1* and *Gad2* expression was widespread in OB neurons supporting their GABAergic identity with weaker expression in NBs (Lledo et al., 2008; Nagayama et al., 2014) (**Figure 1B**).

Next, we surveyed the Allen Mouse Brain Atlas ISH database for subtype-specific markers to examine subtype distribution in the multilayered OB. Using combinations of markers per subtype, we confirmed enrichment of Neuron1, 2, 8 in the granule cell layer, while Neuron4, 5, 6 markers showed enrichment in the glomerular layer (**Figure 1C, Figure S1D**). Neuron7 and 9 markers were predominantly expressed in the anterior olfactory nucleus and external plexiform layer respectively, and Neuron3 was enriched in the mitral cell layer and the external plexiform-glomerular layer boundary (**Figure 1C**). Consistent with scRNA-seq data, immunostainings validated the TdTom co-localization with subtype-specific markers in their predetermined positions, demonstrating the broad repertoire of OB neurons expressing the fate-mapping reporter (**Figure 1D**).

### Dissection of neuronal lineage progression revealed novel gene expression features and key cellular intermediates

To address the complexity of adult neuronal lineage progression and regulation, we generated force-directed graphical visualizations (Weinreb et al., 2017) of neuronal lineage trajectories from V-SVZ astrocyte clusters to OB neurons both in NESFLPO and GCERT2 (12,334 and 7,903 cells, respectively). The resulting trajectories revealed that V-SVZ and OB NBs co-cluster, and lineage progression is constricted by two developmental transitions (**Figure 2A, Table S2**). To identify gene signatures associated with neuronal differentiation stages, we factorized the data with single-cell Hierarchical Poisson Factorization (scHPF) (Levitin et al., 2019), and projected cell scores from different factors onto neuronal lineage trajectories. We identified signatures of two intermediate populations corresponding to factors 2 and 5 (**Figure 2B, Table S2, Figure S2A**). In the OB, migrating NBs with high cell scores for factor 5 were marked by *Prokr2* and *Htr3a*, mutants of which impair NB migration to the OB (Garcia-Gonzalez et al., 2017; Wen et al., 2019). *Notum*, which encodes a secreted WNT inhibitor, was the highest-ranked gene in factor 2 and marked the V-SVZ transition between qNSCs and aNSCs. Factor 2 was also enriched for *Ascl1* and *Sfrp1*, which encodes another secreted WNT inhibitor, and was distinctly separated from factor 1, enriched for astrocyte genes such as *Gfap, Slc1a3*, and *Cldn10* (**Figure 2B**).

**Figure 2.**
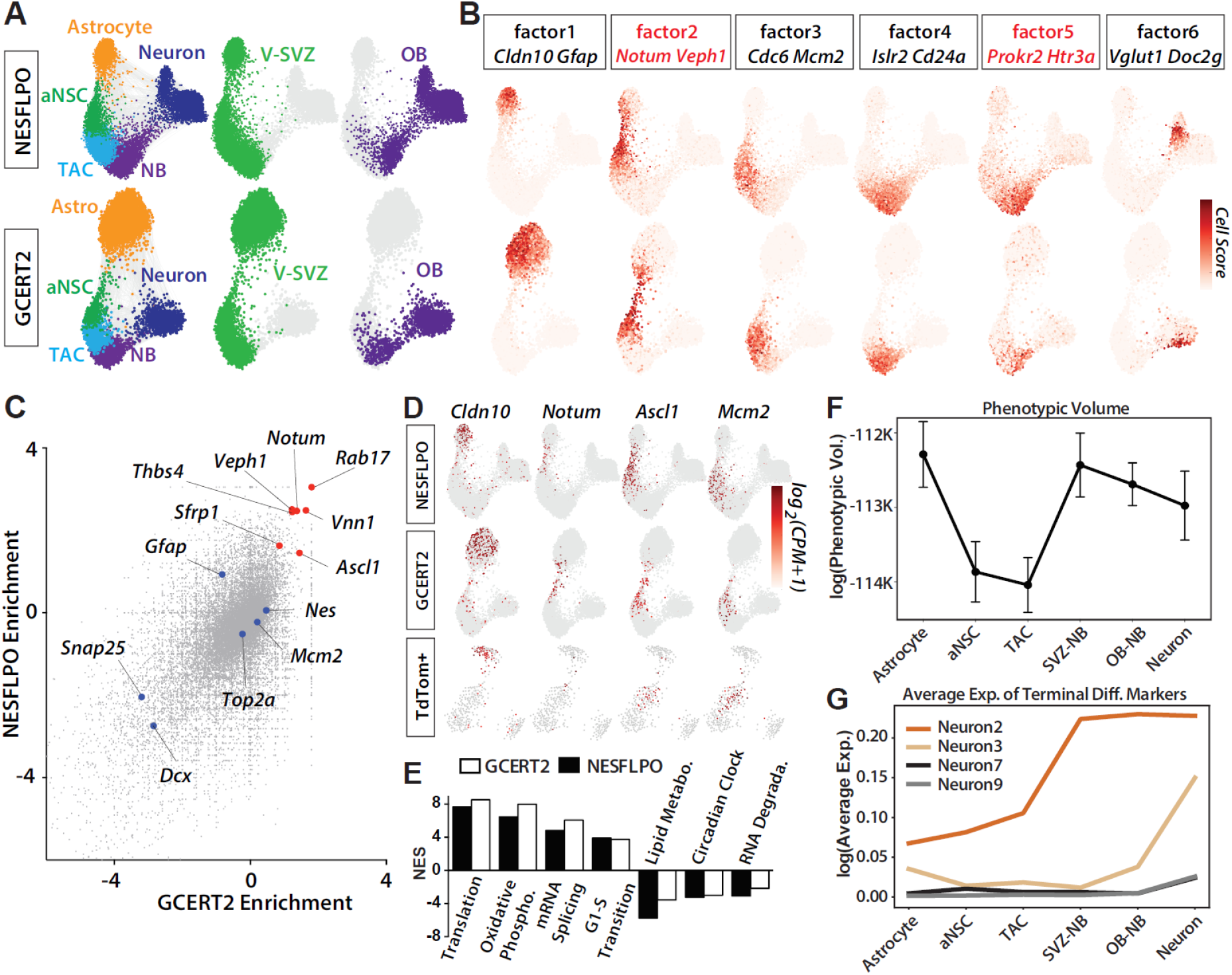
Dissection of neuronal lineage progression reveals an intermediate NSC population marked by *Notum* expression. **(A)** Force-directed neuronal lineage trajectories comprising V-SVZ astrocytes, aNSCs, TACs, and NBs, and OB NBs and neurons both in NESFLPO and GCERT2. Each cell type and/or sample is colored separately. **(B)** Cell scores for six scHPF factors are projected on force-directed lineage trajectories. Two genes with high gene scores are highlighted for each factor. Genes marking the transition states are colored in red. **(C)** GCERT2 and NESFLPO binomial test enrichment of astrocyte and aNSC clusters compared to the remaining neuronal lineage. **(D)** Log_2_(CPM+1) expression values of *Cldn10, Notum, Ascl1* and *Mcm2* in GCERT2, NESFLPO, TdTom+ cells isolated from GCERT2 and NESFLPO. Cells without marker detection are colored in gray. **(E)** Representative enriched and depleted canonical pathways (FDR < 0.01) in *Notum*+ neuronal lineage cells (NES: Normalized enrichment score). **(F)** Phenotypic volume analysis of different clusters in neuronal lineage. **(G)** Average expression of the top five most specific OB neuron subtype markers in different cell clusters based on the binomial test (FDR < 0.01, Table S1).

To highlight markers of the qNSC-aNSC transition, we analyzed genes enriched in astrocyte and aNSC clusters compared to the rest of the lineage, which also revealed *Notum* as a top marker of the intermediate cells (**Figure 2C-D**). *Notum* expression preceded factor 2-enriched *Ascl1* expression in the neuronal lineage suggesting additional heterogeneity (**Figure 2D**). To further investigate the heterogeneity among the intermediate cells, we calculated the pointwise mutual information (PMI), a measure of co-expression, for all gene pairs. As expected, factor 2-enriched genes had high PMIs, but also formed significantly distinct co-expression modules (**Figure S2B**) indicating multistep differentiation in the intermediate population and/or heterogeneity in NSC activation. Importantly, we also observed *Notum* enrichment in V-SVZ endothelial cells (**Figure S2C**) consistent with previous reports of NOTUM expression in cerebral cortex endothelial cells (Zhang et al., 2014). Next, we identified canonical pathways enriched in *Notum*^+^ cells in the neuronal lineage (**Table S3**) by single-cell differential expression (SCDE) and gene set enrichment analysis (GSEA) (Kharchenko et al., 2014; Subramanian et al., 2005). Interestingly, G1/S transition gene set was significantly upregulated further suggesting an advanced cell cycle state (**Figure 2E**).

V-SVZ qNSCs are regionally heterogeneous, generating distinct OB interneuron subtypes depending on their location (reviewed in Chaker et al., 2016; Obernier and Alvarez-Buylla, 2019). We identified several main OB neuron subtypes expressing the fate-mapping reporter; however, it is unknown whether the level of heterogeneity at the gene expression level is preserved along neuronal lineage progression. To formally test this, we calculated the phenotypic volume of each cluster in the lineage trajectory, a metric for cellular heterogeneity (Azizi et al., 2018). Astrocytes formed the most heterogeneous cluster (**Figure 2F**), partly due to the inclusion of qNSCs as well as niche astrocytes in this cluster, while aNSC and TACs were more uniform. We observed an increase in cellular heterogeneity in NBs and OB neurons (**Figure 2F**). An additional non-transcriptional mechanism (e.g. epigenetic memory) may be present in NSCs and TACs to store distinct lineage specification information, and/or the underlying heterogeneity can be masked by cell cycle-related genes. Next, we simply plotted the expression of top markers of neuron subtypes with higher fraction of *TdTom*+ cells (**Table S1, Figure 1B**). Interestingly, transcriptional signatures of neuron subtypes varied in the timing of emergence during the lineage progression. The expression of Neuron2 and 3 markers were enriched in NBs, particularly in OB NBs compared to NSCs and TACs (**Figure 2G**), suggesting an activation of a terminal differentiation program at NB stage, while we did not detect any traces of Neuron9 markers in NBs as expected, as well as the markers of Neuron7 located in anterior olfactory nucleus (**Figure 1B**). Newborn neurons were previously reported in anterior olfactory nucleus (Shapiro et al., 2009) and other olfactory processing areas (Feliciano et al., 2015), however the migration and the maturation status of these cells is not fully clear, and our data suggest a lack of connection between OB NBs and Neuron7.

### SCOPE-seq profiling reveals a dynamic cell size control along neuronal lineage progression and a bias of NOTUM toward proliferating cells

We next sought to determine the spatial arrangement of NOTUM-expressing cells in the V-SVZ. Immunofluorescence revealed strong intracellular NOTUM labeling particularly in the ventral V-SVZ (**Figure 3A, Figure S2D**), consistent with high *Notum* enrichment in that region (Mizrak et al., 2019). Whole-mount staining of the V-SVZ (Mirzadeh et al., 2010; Mirzadeh et al., 2008) showed clear NOTUM labeling on the ventricular surface with partial co-localization between NOTUM and GFAP, which marks V-SVZ qNSCs (**Figure 3B**). To further characterize V-SVZ neuronal lineage cells and the *Notum*^+^ intermediate, we performed SCOPE-seq profiling of >2,500 V-SVZ neuronal lineage cells. SCOPE-seq (Yuan et al., 2018) uses optically decodable mRNA capture beads to link imaging of dissociated single cells (e.g. cell size, surface staining, multiplet detection) captured in microwells with their scRNA-seq profiles (**Figure 3C**). The neuronal lineage trajectory generated from SCOPE-seq recapitulated the lineage progression order as well as *Notum* expression in the intermediate population (**Figure 3C, Table S1**). When optically decoded cells in the neuronal lineage (737/2888 cells) were analyzed (Gunderson et al., 2004; Yuan et al., 2018), we observed a significant increase in cell size at the early aNSC stage (**Figure 3D**). This imaging-based intermediate stage clearly overlapped with *Notum* expressing cells (**Figure 3D**) suggesting *Notum*^+^ cells represent a critical cell size prior to the actively dividing state. Another significant shift in cell size was observed in the early-to-late NB transition, likely due to cell cycle exit. Given the enrichment of G1/S transition gene signatures in *Notum*^+^ cells, the SCOPE-seq data demonstrates a dynamic cell size control during V-SVZ NSC differentiation, where *Notum* marks activating qNSCs at a critical size primed for cell division.

**Figure 3.**
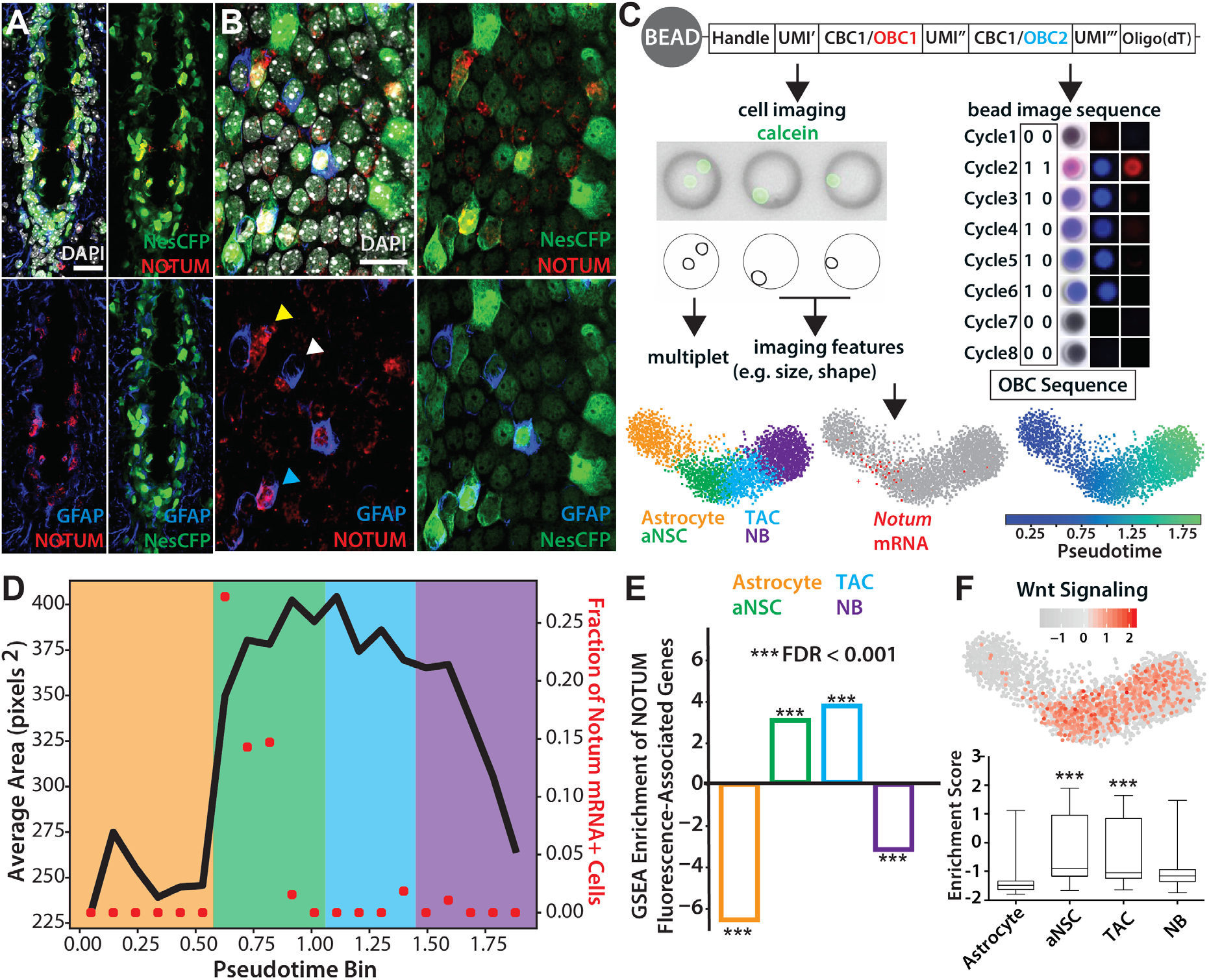
SCOPE-seq profiling of V-SVZ Neuronal Lineage Cells. **(A)** Coronal image of ventral V-SVZ showing NOTUM, NestinCFP (anti-GFP), and GFAP co-localization. Scale bars: 20μm. **(B)** Whole-mount staining showing NOTUM reactivity in ventricle contacting cells on the surface of V-SVZ lateral wall. Cyan arrowhead: NOTUM^+^GFAP^+^, white: NOTUM^−^GFAP^+^, yellow: NOTUM^+^GFAP^−^ cells. Scale bars: 20μm. **(C)** Top: Workflow for SCOPE-seq using optically barcoded mRNA capture beads. Following the barcoded cDNA generation, optical demultiplexing identifies the optical barcode on each bead. Bottom: Pseudotemporal ordering of V-SVZ neuronal lineage cells, and *Notum* marking astrocyte-aNSC transition. Astrocyte: orange, aNSC: green, TAC: blue, NB: purple. **(D)** Average size of pseudotemporally ordered neuronal lineage cells and the fraction of *Notum*+ cells in each pseudotime bin. Shading color corresponds to the stages of differentiation as labeled in **(C)**. **(E)** Enrichment of gene signatures in NOTUM-bound cells in different cell clusters showing significant enrichment in aNSC and TAC. **(F)** GSEA showing the enrichment or depletion of WNT signaling pathway in individual cells, and the quantification of enrichment scores in each cluster.

As a secreted WNT antagonist, NOTUM is active in the extracellular space (Zhang et al., 2015), but can also be retained at the cell surface via binding to glycosaminoglycans on glypicans (**Figure S3A**) (Kakugawa et al., 2015). We used SCOPE-seq to identify NOTUM-bound optically decoded cells (45/737 cells) in the neuronal lineage based on anti-NOTUM surface labeling (**Figure S3B**). Remarkably, these cells showed enrichment of aNSC and TAC cluster gene signatures compared to astrocytes and NBs (**Figure 3E**). Single-cell level GSEA showed significant enrichment of WNT signaling activity in aNSCs and TACs (**Figure 3F**) and depletion in astrocytes, suggesting actively dividing V-SVZ progenitors as potential targets of secreted WNT inhibitors including NOTUM.

### Pharmacological NOTUM inhibition causes aberrant adult neurogenesis

Given these findings and the crucial context-dependent roles of WNT signaling in orchestrating proliferation and differentiation in the V-SVZ (Adachi et al., 2007; Azim et al., 2014; Ortega et al., 2013; Qu et al., 2010; Zywitza et al., 2018), we hypothesized that NOTUM plays a critical role in regulating V-SVZ neuronal progenitors. To examine NOTUM function in the V-SVZ, we blocked NOTUM activity using a selective small molecule inhibitor (ABC99) (Suciu et al., 2018). To investigate effects of NOTUM inhibition *ex vivo* on the V-SVZ, we generated organotypic slice cultures from *Nestin-CFP* reporter mice (Mignone et al., 2004; Wojcinski et al., 2017), and treated them with ABC99 or the inactive control compound ABC101 (**Figure 4A**). After five days of continuous small molecule and EGF treatment in serum free media, we observed a ventral V-SVZ expansion with dense branches of proliferating (EdU^+^) progenitors only in the ABC99 treated slices (**Figure 4B-C, Figure S3C**). Next, ABC99 or ABC101 were intraperitoneally (IP) injected into *Nestin-CFP* mice for three consecutive days to assess the effects of NOTUM inhibition *in vivo* one- and five days post-final injection (1 dpi and 5 dpi) (**Figure 4D**). Systemic ABC99 administration showed no obvious adverse effects on animal health, as previously reported (Pentinmikko et al., 2019). To measure NOTUM activity and to ensure systemic treatment leads to blockage of NOTUM activity in the V-SVZ, we used mass spectrometry-based activity-based protein profiling (MS-ABPP) (reviewed in Cravatt et al., 2008), and developed a targeted LC-MS/MS method to detect NOTUM-derived peptides. Using this technique, we detected significant NOTUM activity in the V-SVZ (**Table S5**), and a 40-50% inhibition of V-SVZ NOTUM after ABC99 injections (**Figure 4E, Figure S3E**). We confirmed WNT pathway activation in the V-SVZ following NOTUM inhibition via qRT-PCR for *Axin2* at 1 dpi (**Figure S3D**) and observed no difference in cell death in the V-SVZ for either ABC99- and ABC101-administered animals. Immunostainings revealed a robust increase in EdU^+^ dividing progenitors as well as Doublecortin-expressing (DCX^+^) NBs in the ABC99-treated animals at 1 dpi (**Figure 4F-G, Figure S3F**). The increase in NBs was much higher in the ventral V-SVZ, where NOTUM expression is enriched (**Figure 3A**), forming pockets of irregular NB expansion (**Figure 4B, 4J**). The effect on NB expansion was transient and disappeared at 5 dpi (**Figure 4H-I**). Consistent with increased NB levels (**Figure 4G, 4J**), we observed a noticeable thickening of the V-SVZ dorsal wedge, which houses many migrating progenitors (**Figure 4B, 4K**). In sum, NOTUM mediated WNT signaling inhibition targets NSC progeny and prevents their over-expansion in the V-SVZ.

**Figure 4.**
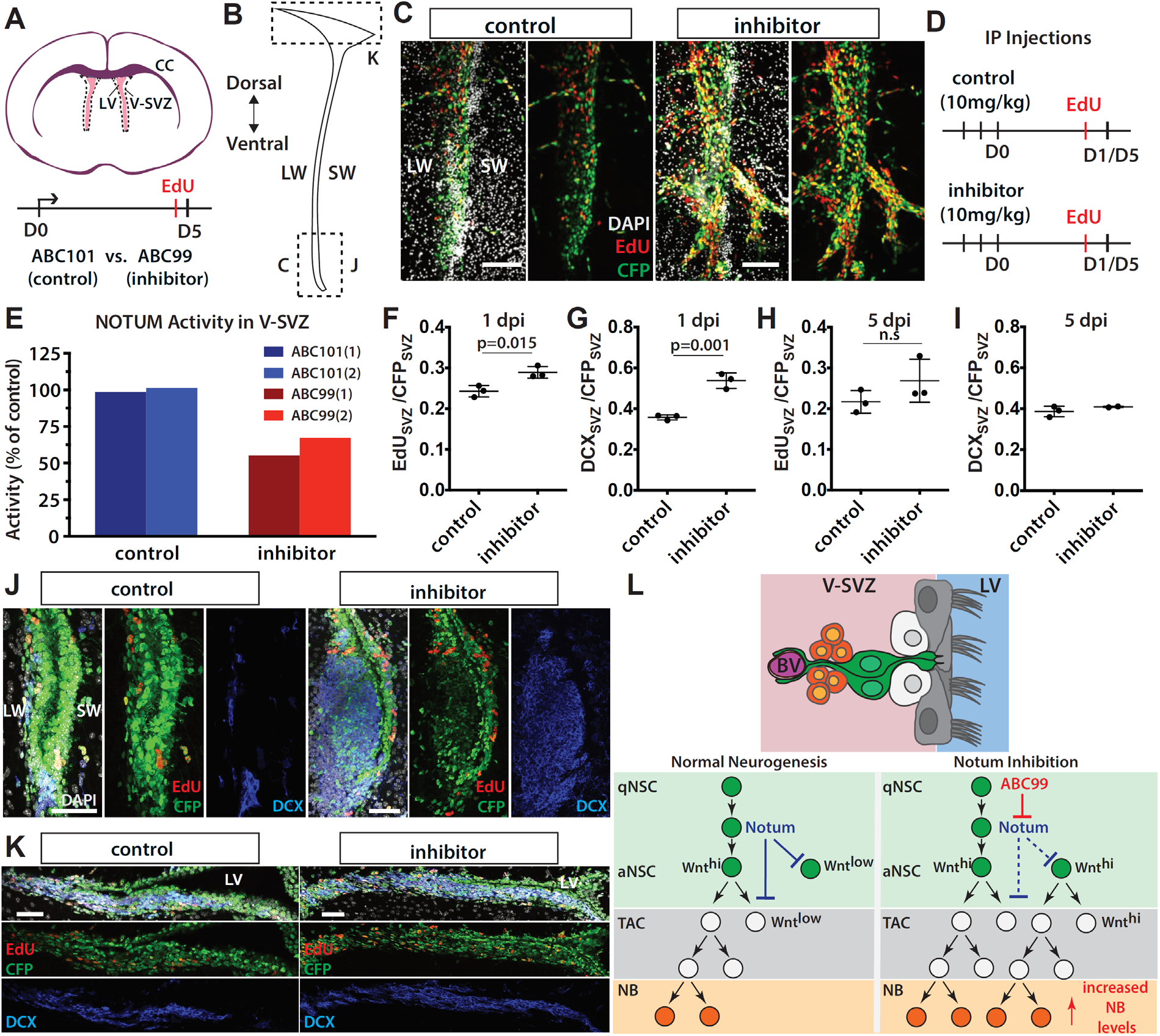
Pharmacological NOTUM inhibition causes aberrant adult neurogenesis. **(A)** Left: Schema of organotypic slice cultures. Slices were treated with ABC101 or ABC99 for five consecutive days (1 μM each). LV: Lateral Ventricle, LW: Lateral Wall, SW: Septal Wall, CC: Corpus Callosum. **(B)** Coronal schema of V-SVZ showing the location of the images in Figs. 4C, 4J-J. **(C)** Rostro-caudal level matched images of the control or the inhibitor treated V-SVZ slices. Scale bars: 100μm. **(D)** Schema showing IP injection regimen. EdU was injected one hour prior to sacrifice. **(E)** Targeted MS-based ABPP showing reduced NOTUM activity in V-SVZ 4 hours after administration of NOTUM inhibitor ABC99 (10 mg/kg, IP) vs. inactive control ABC101. **(F-I)** Quantification of EdU^+^ / CFP^+^ and DCX^+^ / CFP^+^ showing a significant increase in NSC progeny in the inhibitor injected animals (n=3 replicates, mean+/-standard deviation(std)) at 1 dpi and a recovery at 5 dpi. **(J)** Representative rostro-caudal level matched images showing the increase in EdU^+^ and DCX^+^ cells in the ventral V-SVZ in ABC99-treated animals. NestinCFP (anti-GFP) intensity is stronger in ventricle-contacting NSCs, and ependymal cells, and noticeably weaker in NBs indicating the weak CFP perdurance in NSC progeny. **(K)** Representative dorsal wedge images of V-SVZ. **(L)** Schema of the organization of adult V-SVZ NSCs and their progeny, and a mechanism of NOTUM action in adult neurogenesis (BV: blood vessel).

## DISCUSSION

Regulation of adult neural stem cells (NSCs) and their progeny is achieved by stringent molecular checkpoints to prevent excessive activation which could result in stem cell depletion or pathological conditions. The adult V-SVZ is continuously bathed in proliferative WNT ligands from cerebrospinal fluid (Lehtinen et al., 2011) and local production in the niche (**Figure S3G**). Most adult V-SVZ NSCs divide symmetrically to either self-renew or differentiate (Obernier et al., 2018), and as a result are more prone to rapid depletion. Therefore, conservative molecular mechanisms must be in place to cope with the proliferative signals in the niche (e.g. WNT ligands) ensuring high-efficiency neurogenesis. We used scRNA-seq and SCOPE-seq to identify a negative regulator of WNT signaling, *Notum*, as a key marker of poised V-SVZ NSCs. We propose that as poised NSCs become activated, they release NOTUM to laterally inhibit NSC progeny in the niche to prevent their uncontrolled expansion (**Figure 4L**), thus likely providing a more favorable environment for their daughter cells.

Elucidating the factors and signaling mechanisms that mediate the interaction between NSCs and their progeny is essential to develop strategies for mitigating tumor growth or promoting tissue regeneration since pathological conditions drive new V-SVZ derived neurons to migrate to the site of injury (reviewed in Chaker et al., 2016). In this study, we identified an intermediate NSC population from a large-scale scRNA-seq dataset using scHPF (Levitin et al., 2019) to which we attribute a key functional role. Transitioning cells represent the dynamic nature of the V-SVZ niche. Since they are primed to generate multiple progenitors, their communication with the other progenitors in the niche is essential to adapt to impending precursors. Similar strategies may also be employed by stem cell intermediates in other adult tissues and organs ensuring on-demand conservative cell replenishment.

Our analysis also revealed a profound restructuring of cell morphology during lineage progression with several cell size intermediates. Cell cycle and volume need to be tightly coordinated to maintain cellular homeostasis. Particularly, progression through G1/S and G2/M phases has been shown to be size dependent (R. Jones et al., 2017). It will be important to investigate molecular mechanisms underlying the cell size changes during NSC differentiation (e.g. nutritional state, translational activation, circadian clock regulation), whether these changes are reversible in the intermediate populations, and to identify the critical size thresholds NSCs commit to activation or exiting cell cycle. Understanding these mechanisms could offer an artificial means of modulating NSC behavior by regulating their size.

We defined molecularly distinct OB neuron subtypes and validated the repertoire of V-SVZ-derived OB neuron diversity by using different reporter mice combined with scRNA-seq. Progression along V-SVZ to OB neuronal trajectories based on scRNA-seq data demonstrates a convergence of heterogeneous qNSCs to more uniform aNSC and TAC stages, and a late branching during NB to neuron differentiation. These dynamics suggest an additional level of cell-intrinsic programming, such as epigenetic heterogeneity among V-SVZ NSCs and TACs. For example, diverse sets of transcription factors expressed in regionally distinct qNSCs (reviewed in Chaker et al., 2016) may establish epigenetic memory that is conserved in aNSCs and TACs, which later contributes to a later specification event taking place at NB stage. Future single-cell epigenomic profiling studies of V-SVZ NSCs and their progeny could address this complexity. Importantly, our comprehensive V-SVZ-OB dataset provides a resource to map epigenetic changes onto single-cell gene expression data.

Finally, SCOPE-seq enables direct imaging of protein surface binding by imaging-based detection of fluorescently labeled cells and avoids flow sorting which significantly reduces cell quality and changes cellular composition, particularly in the brain. Unlike the DNA-barcoded antibody based approach (Stoeckius et al., 2017), SCOPE-seq can be utilized to detect cells expressing intracellular reporters such as lineage tracing markers commonly employed in stem cell studies while simultaneously providing morphological insight, such as cell size. Developmental trajectories in different germinal regions consist of cells with a broad distribution of cell imaging features (e.g. cell size, cell shape, marker expression); therefore SCOPE-seq based single cell expression resources can be utilized to rapidly identify and quantify different stem cell states including the rare cell populations in a tissue of interest, under various physiological conditions.

## SUPPLEMENTARY FIGURES

**Figure S1.**
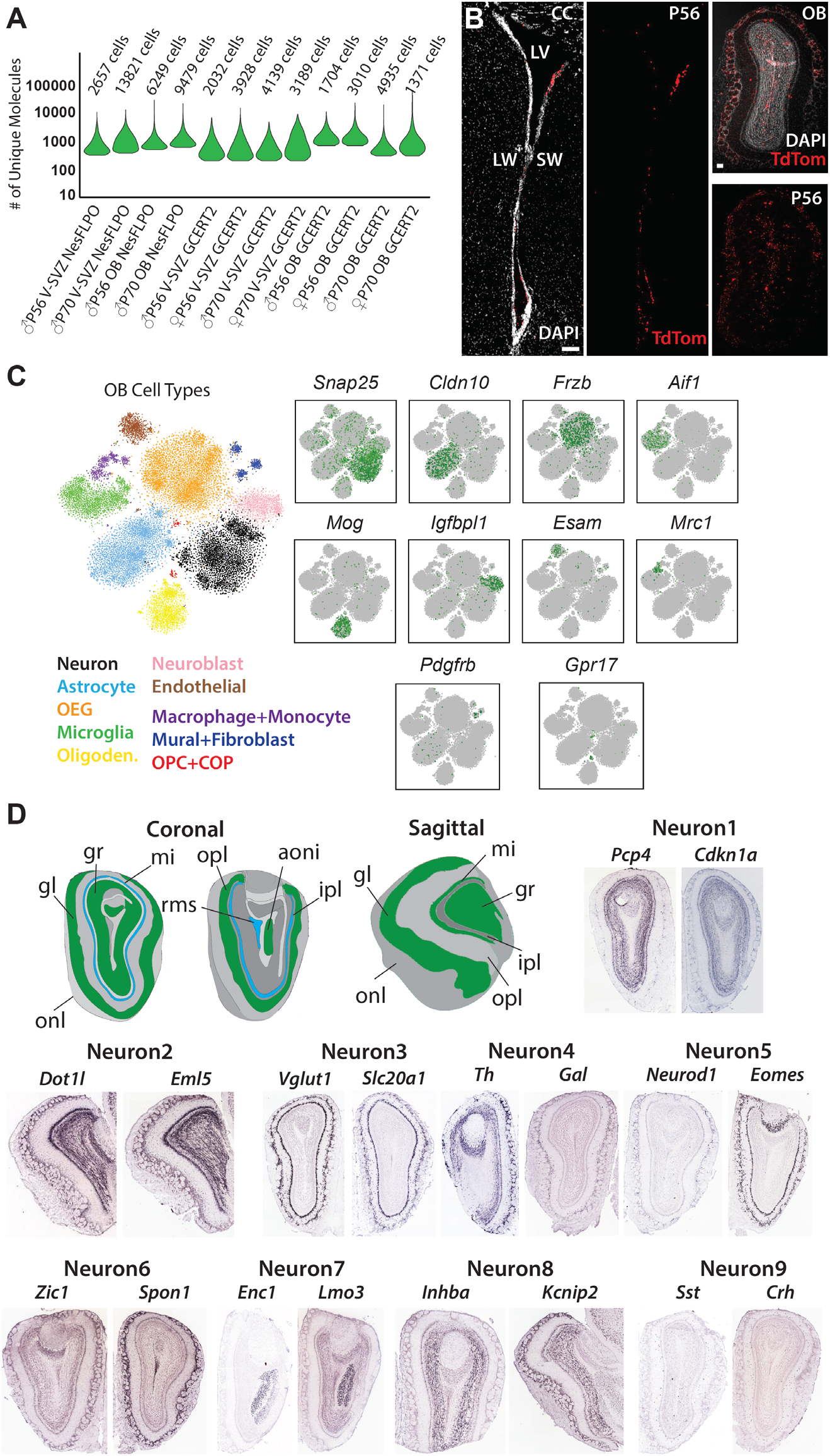
**(A)** Violin plots showing the distribution of unique molecules detected per cell, and the cell numbers in twelve scRNA-seq samples. **(B)** Endogenous TdTom signal in V-SVZ and OB four weeks after tamoxifen injection regimen. TdTom signal is strong even after fixation, and does not require immunostainings. Scale bars 100μm. **(C)** t-SNE projections (Van der Maaten and Hinton, 2008) of major OB cell types colored separately (left) and scaled expression of cluster markers (FDR < 0.001). **(D)** ISH data from Allen Mouse Brain Atlas (Lein et al., 2007) (coronal and sagittal sections) showing the spatial distribution of cluster markers used to determine the location of the neuron subtypes.

**Figure S2.**
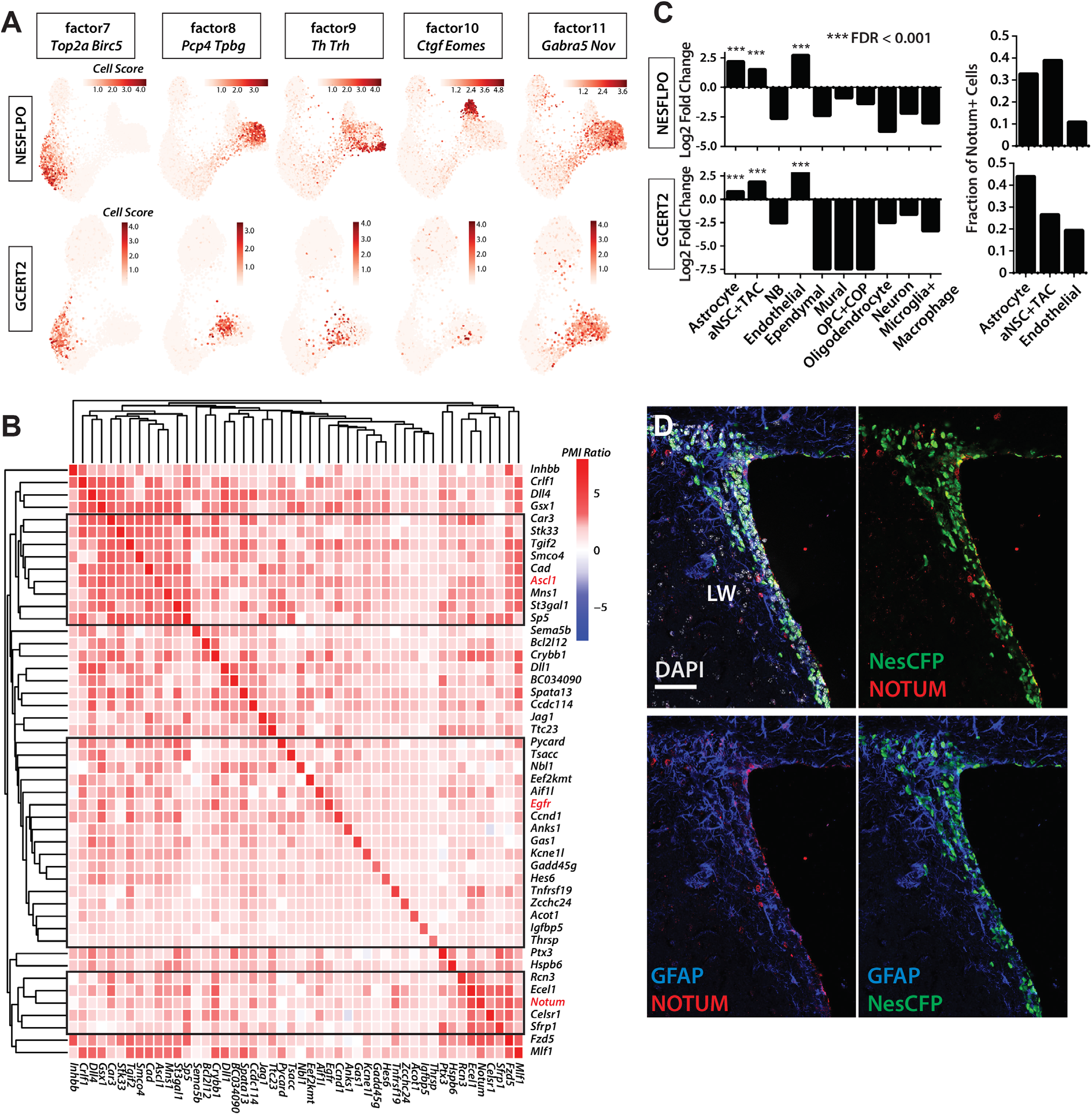
**(A)** Cell scores for the remaining scHPF factors (**Table S2**) are projected on force-directed lineage trajectories. Two genes with high gene scores are highlighted for each factor. **(B)** Heatmap showing the high co-expression probability ratios of top 50 genes (detected in more than 30 cells) in factor 2, and the significantly different clusters formed by them. Three clusters with more than 2 genes were highlighted in black boxes, and a key identifier gene in each cluster was colored in red. Significant clusters were identified after the removal of an outlier cluster formed by *Olig2* and *Hes5*, resulting in 48 genes in the heatmap. **(C)** Left: Column plot showing the significant enrichment of *Notum* expression in V-SVZ Astrocytes, aNSC+TAC and endothelial cell clusters compared to other V-SVZ cells. *Notum* is significantly depleted in NBs and ventricle contacting ependymal cells. Right: The majority of *Notum*+ cells in V-SVZ are in astrocyte, aNSC and TAC clusters. **(D)** Coronal images of NOTUM expression in the dorsal V-SVZ. LW: Lateral Wall. Scale bars 50μm.

**Figure S3.**
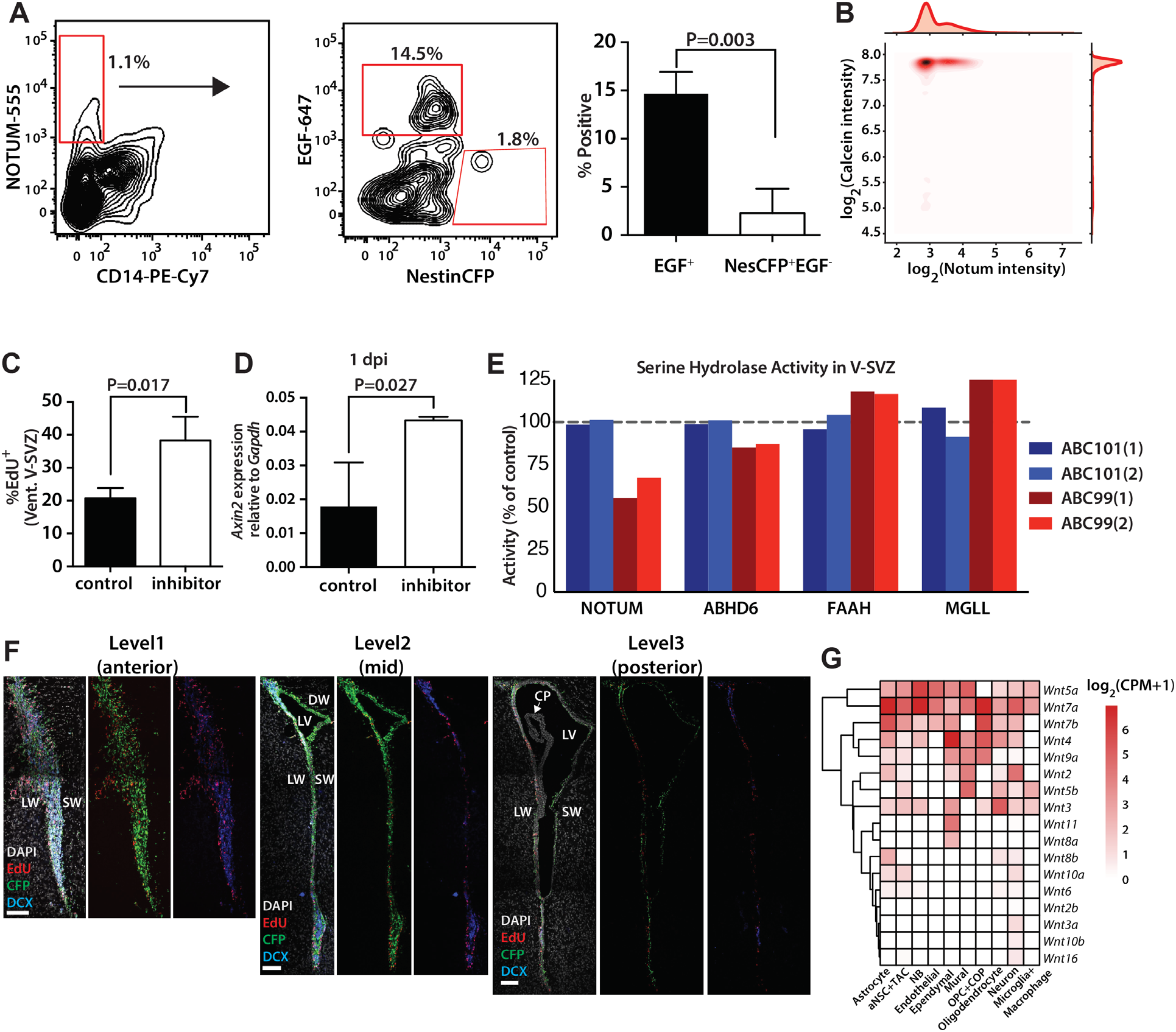
**(A)** Left: Flow cytometric analysis of NOTUM bound cells showing strong co-labeling with EGF as compared to NestinCFP. Right: Quantification of EGF^+^, and NestinCFP^+^EGF^−^ cell percentages among NOTUM^+^ cells (n=3 biological replicates, mean+std). EGF-647 was used to label activated NSC progeny (Codega et al., 2014). **(B)** Log transformed Calcein and Notum staining intensities of SCOPE-seq cells. **(C)** Quantification of EdU^+^ cells in the ventral V-SVZ slice cultures (n=3 sections, mean+std). **(D)** qRT-PCR data indicating *Axin2* upregulation in V-SVZ after the inhibitor injections at 1 dpi (n=3 biological replicates, mean+std). **(E)** Serine hydrolase activity in V-SVZ following ABC101 or ABC99 injections. **(F)** Representative V-SVZ images showing the different rostro-caudal levels analyzed for the quantifications (Level 1: Bregma +1.4mm, Level 2: Bregma +0.9mm, Level 3: Bregma 0.0mm). DW: Dorsal Wall, CP: Choroid Plexus. Scale bars 100μm. **(G)** Heatmap showing the average expression (log_2_(CPM+1)) of WNT ligands in different V-SVZ cell types.

## SUPPLEMENTARY TABLE LEGENDS

**Table S1.** Binomial test results showing the cell type specificity of genes in OB neuron subtypes as well as neuronal lineage cells in GCERT2, NESFLPO and SCOPE-seq datasets. Ensg: Ensemble gene ID. Effect size: log_2_(fraction of cells in cluster with gene detected/fraction of other cells) FDR: False discovery rate. Related to **Figures 1–3, Figure S1**.

**Table S2.** Gene and cell scores of eleven scHPF factors identified in GCERT2 and NESFLPO. Related to **Figure 2** and **Figure S2**.

**Table S3.** Differentially expressed genes in *Notum*+ cells in GCERT2 and NesFLPO, and preranked GSEA results showing the enrichment or depletion of canonical pathways in *Notum*+ cells as compared to the rest of the neuronal lineage cells. Related to **Figure 2**.

**Table S4.** Oligonucleotide sequences on SCOPE-seq mRNA capture beads. Related to **Figure 3**.

**Table S5**. Parallel reaction monitoring results for V-SVZ serine hydrolase activity in ABC101 and ABC99-treated samples. Related to **Figure 4**.

## EXPERIMENTAL PROCEDURES

### Animals

All experiments were performed according to protocols approved by IACUC at the Columbia University and the Memorial Sloan Kettering Cancer Center. Animals were given access to food and water *ad libitum* and were housed on a 12 hr light/dark cycle, and sacrificed at the same time of day. The following mouse lines were used for the experiments: *Nestin-CFP*(Mignone et al., 2004; Wojcinski et al., 2017), B6.Cg-Tg(GFAP-cre/ERT2)505Fmv/J mice (Stock no: 012849, The Jackson Laboratories) (Ganat et al., 2006), *Nestin-FlpoER* (Lao et al., 2012; Wojcinski et al., 2017), *Rosa26^LSL-TdTomato^* (*Ai14*, Stock: 007909, The Jackson Laboratories), *Rosa26^FRT-STOP-FRT-TdTomato^* derived from Ai65 (Stock no: 021875, The Jackson Laboratories), and C57BL/6J (Stock no: 00664, The Jackson Laboratories).

For *TdTomato* induction, four- or six-weeks old mice were injected with 100 mg/kg of tamoxifen (Sigma) intraperitoneally (IP) for three consecutive days, and sacrificed four weeks after the last injection. ABC99 and ABC101 were injected at 10mg/kg for three consecutive days (Related to **Figure 4**). Subcutaneous EdU (50 mg/kg) injections were done one hour before the animals were sacrificed.

### Single cell sequencing sample preparation

Whole-mounts of the lateral and septal walls of the V-SVZ were dissected as previously described (Mirzadeh et al., 2010; Mizrak et al., 2019), and combined prior to dissociation. Olfactory bulb (OB) was also removed to profile V-SVZ and OB from the same animal. For scRNA-seq experiments, sex and age of each animal, and the number of single cells analyzed are indicated in the **Figure S1**. Dissected whole mounts were dissociated as described previously (Mizrak et al., 2019). Briefly, minced V-SVZ pieces were digested with papain (Worthington, 6 mg per sample, 10 min at 37°C) in PIPES solution (120 mM NaCl, 5 mM KCl, 20 mM PIPES (Sigma), 0.45% glucose, 1x Antibiotic/Antimycotic (Gibco), and phenol red (Sigma) in water; pH adjusted to 7.6). After trituration to single cells in DMEM/F12 containing ovomucoid (Worthington, 0.7 mg/ml) and DNAse (Worthington, 0.5 mg/ml), the cell suspension was layered on top of 22% Percoll (Sigma) and centrifuged for 10 mins at 4°C without brakes to remove debris and myelin. The single cell suspension was gently treated with red blood cell lysis buffer (Sigma) for 1 minute, washed by two rounds of centrifugation (1300 rpm at 4°C) and resuspended in 1x Tris Buffered Saline. Cells were stained with Calcein AM live stain dye (Fisher, 1:500) on ice for 30 minutes, and were passed through a 40 μm cell strainer (Fisher) to remove any cell clumps before loading.

### Single cell library preparation and sequencing

Single cell capture and reverse transcription (RT) were performed as previously described (Yuan and Sims, 2016). Cells were loaded in the microwell devices at a concentration of 400-1000 cells per μl. Cell capture, lysis (2-Mercaptoethanol: Fisher Scientific, Buffer TCL: Qiagen), and reverse transcription (Maxima H Minus Reverse Transcriptase: ThermoFisher, SUPERaseIN: Thermofisher, Template switch oligo: IDT) were all performed on 150K microwell devices (microwell size: x: 50 μm, y: 50 μm, z: 58 μm) controlled by an automated microfluidics system. The devices were scanned on a fluorescence microscope (Eclipse Ti-U, Nikon) during the RNA capture step to check for lysis efficiency. RT reactions were performed on Drop-seq beads (MACOSKO-2011-10, ChemGenes) also captured in the microwells. Beads were collected from the devices and pooled for cDNA amplification step. Following exonuclease I treatment (New England Biolabs), cDNA was amplified by PCR for 14 cycles (KAPA HotStart ReadyMix, Kapabiosystems), and used as input for Nextera tagmentation reactions. cDNA and library quality was assessed using both Qubit and Bioanalyzer (Agilent Technologies). All DNA purification steps were carried out using Ampure XP beads (Beckman). High-quality samples were sequenced on a NextSeq 500 sequencer using a 75 cycle High Output Kit (Illumina, 21 cycle read 1, 63 cycle read 2, and 8 cycle index read).

### Single cell sequencing data processing and clustering

The sequencing reads were demultiplexed, aligned, and quantified as described previously (Mizrak et al., 2019). Briefly, read 2 was trimmed to remove 3’ polyA tails (>7 A’s), and fragments with fewer than 24 remaining nucleotides were discarded. Trimmed reads were aligned to the Mus musculus genome and transcriptome annotation (GRCm38, Gencode annotation vM10) using STAR v.2.5.0 with parameters --*sjdbOverhang 65* --*twopassMode Basic* --*outSAMtype BAM Unsorted* (Dobin et al., 2013). Only reads with unique, strand-specific alignments to exons were kept for further analysis. *TdTomato* allele was originally designed as tdTomato::WPRE::polyA. As the sequencing data is 3’ biased, we used the WPRE sequence with flanking sequences to detect *TdTomato* expression in cells. WPRE is denoted as WRPE in the expression matrices.

We extracted 12-nt cell barcodes (CBs) and 8-nt unique molecular identifies (UMIs) from read 1. Degenerate CBs containing either ‘N’s or more than four consecutive ‘G’s were discarded. Synthesis errors, which can result in truncated 11-nt CBs, were corrected similarly to a previously reported method (Shekhar et al., 2016). Briefly, we identified all CBs with at least twenty apparent molecules and for which greater than 90% of UMI-terminal nucleotides were ‘T’. These putative truncated CBs were corrected by removing their last nucleotide. This twelfth nucleotide became the new first nucleotide of corresponding UMIs, which were also trimmed of their last (‘T’) base. We modified our protocol for commercial Drop-seq beads from Chemgenes with a modified structure. A ‘V’ base was added to the end of the UMI upstream of the poly(T) tail for quality-control purposes. Our analytical pipeline was adjusted to recognize truncated CBs by the presence of excessive ‘T’ bases one nucleotide downstream of the position used in the previous pipeline. All reads with the same CB, UMI, and gene mapping were collapsed to represent a single molecule. To correct for sequencing errors in UMI’s, we further collapsed UMI’s that were within Hamming distance 1 of another UMI with the same barcode and gene. To correct for sequencing errors in cell barcodes, we then collapsed CBs that were within Hamming distance one of another barcode, had at least 20 unique UMI-gene pairs a piece, and had at least 75% overlap of their UMI-gene pairs. Finally, we repeated UMI error correction and collapse using the error-corrected CBs. The remaining barcode-UMI-gene triplets were used to generate a digital gene expression matrix. The number of cells included in each sample was determined based on the inflection point in the cumulative histograms for molecules as previously described (Macosko et al., 2015; Yuan and Sims, 2016).

For clustering, we modified our method (Levitin et al., 2019) to select genes detected in fewer cells than expected given their apparent expression level. Briefly, for variable gene selection only, we normalized the molecular counts for each cell to sum to 1. Genes were then ordered by their normalized expression values. For each gene g, we computed a dropout score as follows:

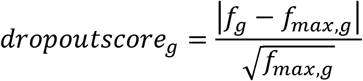

where *f_g_* is the fraction of cells in the dataset that expressed *g*, and *f_max,g_* is the maximum *f_g_* in a rolling window of 25 genes centered on *g*. To cluster and visualize the data, we computed a cell by cell Spearman’s correlation matrix using the marker genes identified above. Using this matrix, we constructed a k-nearest neighbors graph (k=20), which we then used as input to Louvain clustering with Phenograph (Levine et al., 2015). After clustering, we identified genes most specific to each cluster using a binomial test (Shekhar et al., 2016). The same similarity matrix, transformed into a distance matrix by subtracting its values from 1, was used as input to tSNE for visualization. V-SVZ and OB samples from different genotypes (GCERT2 and NESFLPO) were clustered separately. Astrocyte, aNSC, TAC and NB clusters from V-SVZ, and NB and Neuron clusters from OB were merged and re-clustered (k=20), and the resulting k-nearest neighbors graphs were visualized using force-directed graphs to examine their lineage relationship (Weinreb et al., 2017). Doublets, marked by significant enrichment of canonical markers for more than one cell type (e.g. Astrocyte-Oligodendrocytes, Astrocyte-Neuron) were removed from any analysis. Finally, any contaminating cells (e.g. a small cluster of endothelial cells in the larger neuron cluster) formed separate clusters in the sub-clustering analysis, and were removed from the subsequent analyses.

### Single cell hierarchical poisson factorization (scHPF), Pointwise Mutual Information (PMI), phenotypic volume, differential expression and Gene Set Enrichment (GSEA) analyses

scHPF detects latent factors explaining the discrete expression patterns in the scRNAseq datasets (Levitin et al., 2019). Each gene and cell have a score for each factor demonstrating the gene’s contribution to the factor, and the factor’s contribution to the detected expression in the cell respectively. scHPF was applied to the merged neuronal lineage cells from GCERT2 and NESFLPO using mouse protein coding genes. To select the optimal number of factors, first we ran scHPF for different number of factors, *K*. For each value of *K*, we calculated the maximum pairwise overlap of the 100 highest-scoring genes in each factor, and considered overlap significant if p < 0.05 by a hypergeometric test. We picked the model with maximum *K* (p>0.05) and identified the factors with consistent expression patterns both in GCERT2 and NESFLPO, which resulted in eleven factors (**Table S2**).

To assess co-expression and mutual exclusivity of genes, we first identified the significantly expressed genes (detected in more than 30 cells) in the GCERT2 neuronal lineage. For each pair genes, we computed the log-transformed ratio of the joint probability of detecting the two genes in the same cell to the product of their marginal detection probabilities (pointwise mutual information):

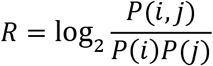

Positive values of *R* indicate co-expression, and negative values indicate mutual exclusivity beyond what would be expected by chance. The results for the top factor 2 genes (**Table S2**) were summarized in **Figure S2B**. Significant clusters of these genes were isolated using R function *pvclust()*, which uses bootstrap resampling to compute p-values (`method.dist=“euclidean”, method.hclust=“average”, nboot=1000; alpha=0.95`).

Phenotypic volume in **Figure 2F** was computed as previously described (Azizi et al., 2018). Briefly, we merged the scRNA-seq count matrices for neuronal lineage cells from the V-SVZ and OB of both the GCERT2 and NESFLPO reporter mice, and applied the Phenograph-based clustering method described above to identify clusters of astrocytes, aNSCs, TACs, neuroblasts, and mature neurons. We transformed the matrix of molecular counts to log_2_(counts per thousand+1), including only genes with a >500 total counts in the dataset. We then computed phenotypic volumes for sub-matrices comprised of astrocyte, aSNC, TAC, SVZ-neuroblasts, OB-neuroblasts, or mature neurons as the product of positive eigenvalues of the gene expression covariance matrix for each of the six populations.

To test the enrichment of WNT signaling pathway (Gene Set M5493-Molecular Signatures Database, Broad Institute) in the SCOPE-seq dataset, we computed the expression of each gene in each cell as the z-scored log_2_(counts per million + 1) and produced a ranked list for each cell. GSEA (GSEA v.2.2.2, http://software.broadinstitute.org/gsea/downloads.jsp) was conducted (Subramanian et al., 2005) for the WNT pathway gene set against each ranked list using the Java implementation using the following command:

~~~
java -cp gsea2-2.2.2.jar -Xmx2512m xtools.gsea.GseaPreranked –scoring scheme_classic –setmin 10 –setmax 1000 –nperm 1000
~~~

Single-cell differential expression (SCDE) analysis was used to identify differentially expressed genes in *Notum*+ cells (Kharchenko et al., 2014). To identify canonical pathways (C2 canonical pathways collection-Broad Institute) enriched or depleted in these cells, SCDE output was pre-ranked based on effect size, and the resulting list was inputted to GSEA using the parameters described above. The results were summarized in **Table S3**.

### Immunohistochemistry, Microscopy, and Image Analysis

8-10 week-old animals were anesthetized, and transcardially perfused with PBS followed by 4% PFA and brains were post-fixed in 4% PFA for 24 hours. After fixation, brains were treated with 30% sucrose until they are embedded in OCT (Tissue-Tek) for cryosectioning. Sections were prepared at 18 μm on cryostat (Leica, CM3050S). Slice cultures were fixed in 4% PFA at end point, and stained at floating sections. Health of the slices were determined based on EdU (Click-IT) incorporation. For immunofluorescent analysis, sections (cryo and slice) were washed with 1X PBS and blocked with blocking buffer (5% BSA in PBS with 0.1% (or 0.3% for slice cultures) Triton-X). Primary antibodies were provided in blocking buffer overnight at 4°C. Washes were performed in PBS with 0.1% Triton-X (or 0.3% for slice cultures). Alexa Fluor conjugated secondary antibodies (Life sciences was supplemented 1/500 in blocking for 1-2 hours at room temperature. After the final washes, EdU staining was performed using Click-IT by manufacturer’s protocol. Hoechst 33258 or DAPI (Invitrogen) was used as counter stain. Slides were mounted using Fluorogel Mounting medium (Electron Microscopy Sciences). The following antibodies were used: rat anti-GFP (1:1000, Nacalai Tesque 04404-84), chicken anti-GFAP (1:1000, Abcam ab4674), rabbit anti-NOTUM (1:500, Sigma abs762), goat anti-DCX (1:500, Santa Cruz sc-8066), chicken anti-TH (1:400, Millipore ab9702), mouse anti-RELN (1:200, Millipore mab5364), rabbit anti-EML5 (1:50, hpa054019), rabbit anti-EOMES (1:500, ab23345), rabbit anti-CALB2 (1:200, Millipore ab5054), and mouse anti-PVALB (1:500, mab1572).

Images were taken using a Zeiss LSM 880 Confocal Microscope or a DM6000 Leica fluorescent microscope. For quantifications, we analyzed sections at three different rostral/caudal levels (from Bregma +1.5 mm to Bregma 0.0 mm) from each replicate (**Figure S3F**). The quantifications from these three levels were averaged to find the total V-SVZ percentages per replicate. Cells in the entire dorsal-ventral extent of V-SVZ were quantified using Fiji software (Schindelin et al., 2012). The identical quantification method was applied to all stainings, replicates and conditions using rostro-caudal level matched sections. Two-way statistical comparisons were conducted using two-tailed unpaired Student’s t-test on three biological replicates, which is similar to sample sizes employed in the field. The quantification data was presented as mean+/-standard deviation.

### Quantitative RT-PCR and Flow Cytometry

ABC99 or ABC101 were injected at 10mg/kg for three consecutive days and the animals were sacrificed at 1 dpi. RNA was isolated from microdissected V-SVZ lateral walls using miRNeasy RNA isolation kit (Qiagen). cDNA was prepared using iScript cDNA preparation kit (BioRad). Following qRT-PCR (Applied BioSystems, Step One) using primers against *Axin2*, relative gene expression levels were quantified using ddCT method. *Gapdh* was used as the housekeeping control to normalize the gene expression. The following primers were used:

*Axin2 F 5’-TGACTCTCCTTCCAGATCCCA-3’*
*Axin2 R 5’-TGCCCACACTAGGCTGACA-3’*
*Gapdh F 5’-CCAAGGTGTCCGTCGTGGATCT-3’*
*Gapdh R 5’-GTTGAAGTCGCAGGAGACAACC-3’*

For flow cytometry, V-SVZ lateral walls from 8-10 weeks old *NestinCFP* animals were dissected, and the single cell suspensions were prepared as described above. Following the Percoll gradient, cells were washed by centrifugation (1300 rpm at 4°C). All staining and washes were performed on ice in 1% BSA, 0.1% Glucose HBSS. After each staining step, cells were washed twice by centrifugation (1300 rpm at 4°C). Cells were first incubated in mouse seroblock FcR (1:100, BioRad buf041A) for 10 minutes to avoid non-specific binding of the antibodies. Cells were then incubated for 15 minutes in anti-NOTUM (1:50), followed by Alexa Fluor 555 secondary antibody staining (1:2000). Finally, cells were stained for 15 minutes with PE-Cy7 conjugated rat anti-CD14 (1:50, Biolegend 123315), and Alexa Fluor 647 conjugated EGF (1:250, Thermofisher E35351). At high concentrations, NOTUM antibody non-specifically interacts with Fc receptors on macrophage, monocyte, microglia, lymphocytes, which can be efficiently blocked with the mouse seroblock FcR incubation (nearly 20-fold depletion in CD14^+^ NOTUM^+^ cells). After the final washes, cells were passed through a 40 μm cell strainer (Fisher) to remove any cell clumps. BD FACS Fortessa was used for flow cytometric analysis, and the data was analyzed using FlowJo. All gates were set using single-color control samples.

### V-SVZ organotypic Slice Cultures

7-10 week old *NestinCFP*/+ animals were euthanized and the brains were dissected into fresh directly into cold HBSS and then embedded in 4% low melting agarose in 1X PBS. Brains were mounted on the chucks and live 250-300um thick coronal sections were obtained using vibratome (Leica). Sections were collected into Neurobasal Media (Invitrogen) supplemented with N2 and B27 Supplements and 20ng/mL EGF (Invitrogen). Sections used for this study spanned the forebrain regions (Bregma +1.4mm to +1mm) Sections were carefully transferred over the membrane inserts (Millipore Millicell cell culture inserts) that are placed over 1mL of media. Alternating sections were treated either with ABC99 or ABC101. Media was changed every day, and 10 ul of media was dropped on each slice to keep them moist. Slices were cultured at 37°C with 5% CO2. Prior to fixation at end point, cultures were pulsed with EdU (2.5μg/mL) for 5 hours.

### SCOPE-seq sample and library preparation, and sequencing

Single cell suspensions of V-SVZ cells from male C57BL/6J animals were prepared as described above. Following the Percoll gradient, cells were washed in 1% BSA, 0.1% Glucose HBSS by centrifugation (1300 rpm at 4°C). Cells were first incubated in anti-NOTUM (1:50) for 15 minutes on ice, washed twice by centrifugation (1300 rpm at 4°C), and incubated in Alexa Fluor 647 secondary antibody (1:2000) for 15 minutes on ice. Cells were then washed twice, and stained with Calcein AM live stain dye (Fisher, 1:500) in 1X TBS on ice for 30 minutes. The resulting cell suspension was passed through a 40 μm cell strainer, and loaded in a microwell device (>130K microwells) at a concentration of 1000 cells per μl. Untrapped cells were flushed out with 1x TBS, and the device was scanned under an inverted automated Nikon Eclipse Ti2 microscope in two fluorescence channels and the bright field channel in ~35 minutes. Optically barcoded (OBC) beads (Chemgenes) were then loaded in the microwell device, and untrapped beads were flushed out with 1x TBS. The microwell device containing the cells and the beads was connected to the computer-controlled reagent and temperature delivery system to perform on-chip RT as described above. The device was scanned during RNA capture to assess the lysis efficiency. After the post-RT washes were completed, the device was disconnected from the automated reagent delivery system. On-chip Exo I treatment was performed at 37°C for 45 minutes followed by TE/TW buffer wash (10 mM Tris pH 8.0, 1 mM EDTA, 0.01% Tween-20). The device was then connected to the automated optical demultiplexing system for reagent delivery.

Optical demultiplexing was performed as previously described with several modifications (Yuan et al., 2018). Each optically decodable mRNA capture bead is conjugated to an oligonucleotide with two, 8-base sequences, each of which is selected from a different set of 96 sequences (**Table S4**). The barcodes are synthesized combinatorially using solid-phase oligonucleotide synthesis (Chemgenes). Together, the two sequences form the cell-identifying barcode for a given bead, and the complete set of cell-identifying barcodes includes 96×96=9,216 possible barcodes. Further multiplexing capacity is achieved by preparing sequencing libraries separately from beads deposited in different sub-arrays of the same device (Yuan et al., 2018). The oligonucleotides also contain interspersed random sequences that form a unique molecular identifier (UMI) and are terminated with poly(dT) to facilitate mRNA capture by hybridization to the poly(A) tail.

To identify the two sequences on each bead that form the cell-identifying barcode, we use sequential fluorescence hybridization. In each cycle of fluorescence hybridization, we introduce a set of 8-base, Cy5-labeled oligonucleotides to probe the first sequence on each bead and a set of 8-base, Cy3-labeled oligonucleotides to probe the second sequence. The probes are pooled such that all 96 sequences in each set can be identified with eight cycles of sequential fluorescence hybridization (Gunderson et al., 2004) (**Table S4**). Each cycle consists of a background scan and probe scan (two fluorescence channels; Cy3 and Cy5, and bright field), and probe hybridization and stripping (2x SSC buffer with tween-20 and 150mM NaOH melting solution). Background scans showed near absolute probe stripping efficiency. Each pool of hybridization mixture consists of Cy3- and Cy5-labeled oligonucleotides complementary to OBCs (96×96 possible combinations). After the optical demultiplexing workflow, all wells were sealed with perfluorinated oil (Sigma, F3556), the device with separated regions was cut in to 15 pieces with razor blades. Beads from each region were placed in a microcentruge tube containing 100% ethanol, and collected by vortexing, sonication, and centrifugation. By preparing and indexing libraries separately from 15 different device regions, we effectively expand our barcoding capacity to 96×96×15 = 138,240 barcodes. The beads were then washed with TE/SDS (10 mM Tris-HCl, 1 mM EDTA, 0.5% SDS) once, TE/TW twice, and nuclease-free water. cDNA was amplified by PCR for 16 cycles (KAPA HotStart ReadyMix, Kapabiosystems), and used as input for Nextera tagmentation reactions. Library preparation was performed as described above resulting in 15 sequencing libraries. These libraries were pooled, and sequenced on a NextSeq 500 sequencer (Illumina) using a 75 cycle High Output Kit (26 cycle read 1, 58 cycle read 2, and 8 cycle index read).

### SCOPE-seq imaging and sequencing data analysis

Image analysis was performed using ImageJ. To identify the OBC sequence, each bead was identified in the bright field channel (`setAutoThreshold (“Default”); setOption (“BlackBackground”, false); run(“Convert to Mask”)`), and Cy3 and Cy5 channel intensities were measured in these regions of interest (ROIs). OBC calling was performed using a bead-by-bead algorithm (Gunderson et al., 2004). Cell ROIs were extracted from the cell loading scan using the calcein signal, the intensity measurements were performed for two fluorescence channels (calcein AM and anti-NOTUM fluorescence). The coordinates for the center of the wells and the well ROIs were extracted from the bright field channel (`Auto Local Threshold, method=Bernsen, radius=30, parameter_1 = 40, parameter_2 = 0`). Cells were assigned to the center of their wells, the wells containing more than one cell were marked as multiplets.

Both the calcein AM and anti-NOTUM fluorescence intensity distributions were bimodal for microwells containing individual cells (**Figure S3B**). We thresholded the anti-NOTUM and calcein AM fluorescence at the intensity between the two modes of each intensity histogram with the fewest cells to identify a set calcein AM^+^/NOTUM^+^ and calcein AM^+^/NOTUM^−^ cells for further analysis. We discarded cells for which the same cell-identifying barcode was associated with more than one cell in the same sub-array of the device.

To analyze the scRNA-seq data from SCOPE-Seq, we first extract the cell-identifying barcode and UMI from Read 1. The sequences in the two sets that are combined to form each cell-identifying barcode have a Hamming distance of at least three for all sequence pairs within each set. Therefore, we correct at most one substitution error in each of the two sequences should one arise from sequencing, oligonucleotide synthesis, or other sources. We only keep reads with a complete cell-identifying barcode. Next, we align the reads from Read 2 using the STAR aligner and keep strand-specific alignments to exons as described above after removing poly(A) tails >7 bases long from the 3’-end of each read. Finally, we use the UMIs to estimate the number of molecules associated with each gene in each cell as described above. Clustering and visualization of the scRNA-seq profiles resulting from SCOPE-seq was carried out as described above for conventional scRNA-seq.

To associate imaging features with pseudotime in the neuronal lineage trajectory, we first extracted neuronal lineage cells from the SCOPE-seq data based on Phenograph clustering of the entire data set. This included all clusters enriched in markers of astrocytes, aNSCs, TACs, and NBs. After marker selection using the drop-out curve for the neuronal lineage cells as described above, we computed a similarity matrix of pairwise Spearman correlation coefficients from which we constructed a k-nearest neighbors graph (k=20) which we projected in two dimensions as a force-directed graph using the *spring_layout* function in the *networkx* Python module. We then fit a polynomial to the resulting two-dimensional projection, which we interpret as the average trajectory. We then assign each cell to a position along this average trajectory by determining its minimum distance from the polynomial curve. Finally, we grouped the cells into 20 bins of equal size along the polynomial trajectory to produce **Figure 3D** where we plot the average area (based on calcein AM fluorescence) of the cells in each bin and the fraction of *Notum* mRNA+ cells in each bin.

To associate NOTUM surface staining with gene expression signatures of the four major subpopulations in the neuronal lineage (astrocyte, aNSC, TAC, and NB), we first computed the pointwise mutual information (PMI) between NOTUM surface staining from SCOPE-seq imaging (a categorical where a cell is either positive or negative) and expression of each gene from scRNA-seq (a second categorical where a gene is either detected or not in a given cell):

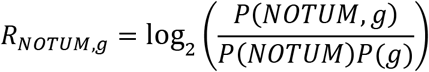

Here, *P*(*NOTUM,g*) is the joint probability that a cell is both positive for NOTUM surface staining and positive for detection of the mRNA encoding gene *g, P*(*NOTUM*) is the probability that a cell is positive for NOTUM surface staining, and *P*(*g*) is the probability that a cell is positive for detection of the mRNA encoding gene *g*. Therefore, if a gene co-occurs with NOTUM surface staining more than expected by chance, *R_NOTUM,g_* is positive. To identify genes that were significantly associated with NOTUM surface staining, we used a permutation test, shuffling the identities of NOTUM surface positive cells 1,000 times and correcting the resulting p-values for multiple comparisons using the Benjamini-Hochberg procedure as implemented in the Python module *statsmodels*. We created a gene set comprised of genes detected in greater than 20 cells with p_corr_ < 0.05. We then created a ranked list of genes for each of the four neuronal lineage clusters using −/+*log*(*p*) where *p* is the p-value from the binomial test used to identify genes specific to each cluster that is described above. This parameter is positive for genes that are positively enriched in a given cluster and negative for genes that are depleted. Finally, we used GSEA to compute normalized enrichment scores for the gene set comprised of NOTUM surface staining-associated genes in the ranked list for each of the four clusters using the GSEA parameters described above. The results are summarized in **Figure 3E**.

### Mass Spectrometry Based Activity-Based Protein Profiling

To prepare V-SVZ proteome, ABC99 or ABC101 were injected at 10mg/kg for three consecutive days and the animals were sacrificed 4 hours after the last injection. Whole-mounts of the lateral wall of V-SVZ were dissected (Mirzadeh et al., 2010) and snap frozen. Frozen samples from each mouse were homogenized using probe sonication in 300 μL of cold DPBS. Protein concentrations were determined using Bio-Rad DC protein assay. Due to the low protein yields, four SVZ proteomes from each group (ABC101 or ABC99-treated) were combined into a single sample and split into two 1.0 mL samples at a protein concentration of 2.7 mg/mL. Tissue homogenates (1.0 mL, 2.7 mg/mL) were treated with FP-biotin (10 μL, 1 mM in DMSO) to a final concentration of 10 μM and incubated for 1 hour at 37°C with gentle rotation. To remove unreacted probe, protein was precipitated by addition of a cold acetone:methanol (9:1) mixture (8 mL) and incubated at −20°C freezer overnight. The samples were centrifuged at 4,700 x *g* for 15 min at 4°C resulting in the formation of a protein pellet at the bottom of the tube. The supernatant was removed by inversion and the pellet was allowed to air dry for 10 min at room temperature. The protein pellet was dissolved in 0.2 mL of a denaturing solution containing urea (8.0 M), sodium dodecyl sulfate (SDS) (1%), and 1,4-dithiothreitol (DTT) (200 mM) in DPBS. Following complete dissolution, the samples were incubated for 30 min at 37°C and then treated with iodoacetamide (20 μL, 500 mM in DPBS) for an additional 30 min at room temperature. The samples were diluted with 0.25% SDS in DPBS (1.4 mL) and incubated with DPBS-washed streptavidin agarose beads (60 μL) for 1 h at room temperature. The streptavidin beads were then sequentially washed with 0.25% SDS (10 x 0.8 mL), DPBS (10 x 1.1 mL), and distilled water (10 x 1.1 mL). The washed beads were transferred to a clean vessel and trypsin (1.6 μg in 100 μl of 2 M urea in DPBS) was added. Samples were incubated overnight at room temperature with agitation. The beads were removed from the digestion using filtration and washed with DPBS (100 μL). The combined filtrates were then acidified by addition of formic acid (20 μL of 100% formic acid) and desalted using SOLAμ™ SPE Plates (HRP 2 mg / 1 mL). Samples were dried by centrifugal evaporation and stored at −80 °C until analysis.

Dry peptide samples were reconstituted in water containing 0.1% formic acid (20 μL) and 10 μL were injected onto an EASY-Spray column (15 cm x 75 μm ID, PepMap C18, 3 μm particles, 100 Å pore size, Thermo Fisher Scientific) using a Dionex RSLCnano LC (Thermo Fisher Scientific). Peptides were separated over a 30 min gradient of 0 to 40% acetonitrile (0.1% formic acid) and analyzed on an Orbitrap Fusion Lumos (Thermo Fisher Scientific) operated using a parallel reaction monitoring (PRM) method for three distinct NOTUM peptides and a single peptide from three other common serine hydrolase targets: ABHD6, FAAH and MGLL. Selected ions were isolated and fragmented by high energy collision dissociation (HCD) and fragments were detected in the Orbitrap at 15,000 or 30,000 resolution. Sequences, targets and targeting parameters can be found in **Table S5** (PRM Parameters).

Raw data files were uploaded analyzed in Skyline (v4.2.0.19009) to determine the relative abundance of each serine hydrolase peptide in vehicle-treated samples relative to ABC99-treated samples. Peptide quantification was performed by calculating the sum of the peak areas corresponding to 6 fragment ions from each peptide (see table below). To determine protein activity relative to the controls, an area ratio was calculated for each fragment ion, dividing the peak area of an ABC99-treated sample by the mean peak area of the ABC101-control samples. The relative protein activity was found from the slope of the regression line obtained by plotting the treated ratios against mean control ratios. The peptides and fragment ions were pre-selected from in-house reference spectral libraries acquired in data-dependent acquisition mode to identify authentic spectra for each peptide. Product ions used for Skyline quantification appear in **Table S5**.

## Acknowledgements

A.L.J. was supported by R01NS092096 from NIH/NINDS. N.S.B. was supported by postdoctoral fellowship C32599GG from NYSTEM. P.A.S. was funded by R33CA202827 from NIH/NCI and R44HG010003 from NIH/NHGRI.

## Author Contributions

D.M. and P.A.S. designed the study. D.M. performed the scRNA-seq and SCOPE-seq experiments. D.M. and P.A.S. analyzed the data. D.M. and N.S.B. performed the immunostainings and the functional validation experiments. J.Y., Z.L., and P.A.S. developed SCOPE-seq. R.S., M.J.N., N.N., K.M.L and B.F.C. performed and analyzed MS-ABPP experiments. D.M., N.S.B., M.J.N., A.L.J., and P.A.S. wrote the manuscript.

## Data Availability

The sequencing data and count matrices have been deposited with the Gene Expression Omnibus (GEO) under accession code GSE134918.

## Code Availability

Data analysis code used for scRNA-seq is available at https://github.com/simslab.

## REFERENCES

Adachi, K., Mirzadeh, Z., Sakaguchi, M., Yamashita, T., Nikolcheva, T., Gotoh, Y., Peltz, G., Gong, L., Kawase, T., Alvarez-Buylla, A., et al. (2007). Beta-catenin signaling promotes proliferation of progenitor cells in the adult mouse subventricular zone. Stem Cells 25, 2827–2836.

Alfonso, J., Le Magueresse, C., Zuccotti, A., Khodosevich, K., and Monyer, H. (2012). Diazepam binding inhibitor promotes progenitor proliferation in the postnatal SVZ by reducing GABA signaling. Cell Stem Cell 10, 76–87.

Azim, K., Fischer, B., Hurtado-Chong, A., Draganova, K., Cantu, C., Zemke, M., Sommer, L., Butt, A., and Raineteau, O. (2014). Persistent Wnt/beta-catenin signaling determines dorsalization of the postnatal subventricular zone and neural stem cell specification into oligodendrocytes and glutamatergic neurons. Stem Cells 32, 1301–1312.

Azizi, E., Carr, A.J., Plitas, G., Cornish, A.E., Konopacki, C., Prabhakaran, S., Nainys, J., Wu, K., Kiseliovas, V., Setty, M., et al. (2018). Single-Cell Map of Diverse Immune Phenotypes in the Breast Tumor Microenvironment. Cell 174, 1293–1308 e1236.

Brill, M.S., Ninkovic, J., Winpenny, E., Hodge, R.D., Ozen, I., Yang, R., Lepier, A., Gascon, S., Erdelyi, F., Szabo, G., et al. (2009). Adult generation of glutamatergic olfactory bulb interneurons. Nat Neurosci 12, 1524–1533.

Chaker, Z., Codega, P., and Doetsch, F. (2016). A mosaic world: puzzles revealed by adult neural stem cell heterogeneity. Wiley Interdiscip Rev Dev Biol 5, 640–658.

Codega, P., Silva-Vargas, V., Paul, A., Maldonado-Soto, A.R., Deleo, A.M., Pastrana, E., and Doetsch, F. (2014). Prospective identification and purification of quiescent adult neural stem cells from their in vivo niche. Neuron 82, 545–559.

Cravatt, B.F., Wright, A.T., and Kozarich, J.W. (2008). Activity-based protein profiling: from enzyme chemistry to proteomic chemistry. Annu Rev Biochem 77, 383–414.

Delgado, R.N., and Lim, D.A. (2015). Embryonic Nkx2.1-expressing neural precursor cells contribute to the regional heterogeneity of adult V-SVZ neural stem cells. Dev Biol 407, 265–274.

Dobin, A., Davis, C.A., Schlesinger, F., Drenkow, J., Zaleski, C., Jha, S., Batut, P., Chaisson, M., and Gingeras, T.R. (2013). STAR: ultrafast universal RNA-seq aligner. Bioinformatics (Oxford, England) 29, 15–21.

Doetsch, F., Caille, I., Lim, D.A., Garcia-Verdugo, J.M., and Alvarez-Buylla, A. (1999). Subventricular zone astrocytes are neural stem cells in the adult mammalian brain. Cell 97, 703–716.

Driehuis, E., and Clevers, H. (2017). WNT signalling events near the cell membrane and their pharmacological targeting for the treatment of cancer. Br J Pharmacol 174, 4547–4563.

Eng, C.L., Lawson, M., Zhu, Q., Dries, R., Koulena, N., Takei, Y., Yun, J., Cronin, C., Karp, C., Yuan, G.C., et al. (2019). Transcriptome-scale super-resolved imaging in tissues by RNA seqFISH. Nature 568, 235–239.

Feliciano, D.M., Bordey, A., and Bonfanti, L. (2015). Noncanonical Sites of Adult Neurogenesis in the Mammalian Brain. Cold Spring Harb Perspect Biol 7, a018846.

Fiorelli, R., Azim, K., Fischer, B., and Raineteau, O. (2015). Adding a spatial dimension to postnatal ventricular-subventricular zone neurogenesis. Development 142, 2109–2120.

Fuentealba, L.C., Rompani, S.B., Parraguez, J.I., Obernier, K., Romero, R., Cepko, C.L., and Alvarez-Buylla, A. (2015). Embryonic Origin of Postnatal Neural Stem Cells. Cell 161, 1644–1655.

Ganat, Y.M., Silbereis, J., Cave, C., Ngu, H., Anderson, G.M., Ohkubo, Y., Ment, L.R., and Vaccarino, F.M. (2006). Early postnatal astroglial cells produce multilineage precursors and neural stem cells in vivo. J Neurosci 26, 8609–8621.

Garcia-Gonzalez, D., Khodosevich, K., Watanabe, Y., Rollenhagen, A., Lubke, J.H.R., and Monyer, H. (2017). Serotonergic Projections Govern Postnatal Neuroblast Migration. Neuron 94, 534–549 e539.

Gunderson, K.L., Kruglyak, S., Graige, M.S., Garcia, F., Kermani, B.G., Zhao, C., Che, D., Dickinson, T., Wickham, E., Bierle, J., et al. (2004). Decoding randomly ordered DNA arrays. Genome Res 14, 870–877.

Kakugawa, S., Langton, P.F., Zebisch, M., Howell, S., Chang, T.H., Liu, Y., Feizi, T., Bineva, G., O’Reilly, N., Snijders, A.P., et al. (2015). Notum deacylates Wnt proteins to suppress signalling activity. Nature 519, 187–192.

Kawaguchi, D., Furutachi, S., Kawai, H., Hozumi, K., and Gotoh, Y. (2013). Dll1 maintains quiescence of adult neural stem cells and segregates asymmetrically during mitosis. Nat Commun 4, 1880.

Kharchenko, P.V., Silberstein, L., and Scadden, D.T. (2014). Bayesian approach to single-cell differential expression analysis. Nature methods 11, 740–742.

Klein, A.M., Mazutis, L., Akartuna, I., Tallapragada, N., Veres, A., Li, V., Peshkin, L., Weitz, D.A., and Kirschner, M.W. (2015). Droplet barcoding for single-cell transcriptomics applied to embryonic stem cells. Cell 161, 1187–1201.

Kohwi, M., Osumi, N., Rubenstein, J.L., and Alvarez-Buylla, A. (2005). Pax6 is required for making specific subpopulations of granule and periglomerular neurons in the olfactory bulb. J Neurosci 25, 6997–7003.

Kriegstein, A., and Alvarez-Buylla, A. (2009). The glial nature of embryonic and adult neural stem cells. Annu Rev Neurosci 32, 149–184.

Lao, Z., Raju, G.P., Bai, C.B., and Joyner, A.L. (2012). MASTR: a technique for mosaic mutant analysis with spatial and temporal control of recombination using conditional floxed alleles in mice. Cell Rep 2, 386–396.

Lehtinen, M.K., Zappaterra, M.W., Chen, X., Yang, Y.J., Hill, A.D., Lun, M., Maynard, T., Gonzalez, D., Kim, S., Ye, P., et al. (2011). The cerebrospinal fluid provides a proliferative niche for neural progenitor cells. Neuron 69, 893–905.

Lein, E.S., Hawrylycz, M.J., Ao, N., Ayres, M., Bensinger, A., Bernard, A., Boe, A.F., Boguski, M.S., Brockway, K.S., Byrnes, E.J., et al. (2007). Genome-wide atlas of gene expression in the adult mouse brain. Nature 445, 168–176.

Levine, J.H., Simonds, E.F., Bendall, S.C., Davis, K.L., Amir el, A.D., Tadmor, M.D., Litvin, O., Fienberg, H.G., Jager, A., Zunder, E.R., et al. (2015). Data-Driven Phenotypic Dissection of AML Reveals Progenitor-like Cells that Correlate with Prognosis. Cell 162, 184–197.

Levitin, H.M., Yuan, J., Cheng, Y.L., Ruiz, F.J., Bush, E.C., Bruce, J.N., Canoll, P., Iavarone, A., Lasorella, A., Blei, D.M., et al. (2019). De novo gene signature identification from single-cell RNA-seq with hierarchical Poisson factorization. Mol Syst Biol 15, e8557.

Liu, X., Wang, Q., Haydar, T.F., and Bordey, A. (2005). Nonsynaptic GABA signaling in postnatal subventricular zone controls proliferation of GFAP-expressing progenitors. Nat Neurosci 8, 1179–1187.

Lledo, P.M., Merkle, F.T., and Alvarez-Buylla, A. (2008). Origin and function of olfactory bulb interneuron diversity. Trends Neurosci 31, 392–400.

Lopez-Juarez, A., Howard, J., Ullom, K., Howard, L., Grande, A., Pardo, A., Waclaw, R., Sun, Y.Y., Yang, D., Kuan, C.Y., et al. (2013). Gsx2 controls region-specific activation of neural stem cells and injury-induced neurogenesis in the adult subventricular zone. Genes Dev 27, 1272–1287.

Macosko, E.Z., Basu, A., Satija, R., Nemesh, J., Shekhar, K., Goldman, M., Tirosh, I., Bialas, A.R., Kamitaki, N., Martersteck, E.M., et al. (2015). Highly Parallel Genome-wide Expression Profiling of Individual Cells Using Nanoliter Droplets. Cell 161, 1202–1214.

Merkle, F.T., Fuentealba, L.C., Sanders, T.A., Magno, L., Kessaris, N., and Alvarez-Buylla, A. (2014). Adult neural stem cells in distinct microdomains generate previously unknown interneuron types. Nat Neurosci 17, 207–214.

Merkle, F.T., Mirzadeh, Z., and Alvarez-Buylla, A. (2007). Mosaic organization of neural stem cells in the adult brain. Science 317, 381–384.

Mich, J.K., Signer, R.A., Nakada, D., Pineda, A., Burgess, R.J., Vue, T.Y., Johnson, J.E., and Morrison, S.J. (2014). Prospective identification of functionally distinct stem cells and neurosphere-initiating cells in adult mouse forebrain. Elife 3, e02669.

Mignone, J.L., Kukekov, V., Chiang, A.S., Steindler, D., and Enikolopov, G. (2004). Neural stem and progenitor cells in nestin-GFP transgenic mice. J Comp Neurol 469, 311–324.

Mirzadeh, Z., Doetsch, F., Sawamoto, K., Wichterle, H., and Alvarez-Buylla, A. (2010). The subventricular zone en-face: wholemount staining and ependymal flow. J Vis Exp.

Mirzadeh, Z., Merkle, F.T., Soriano-Navarro, M., Garcia-Verdugo, J.M., and Alvarez-Buylla, A. (2008). Neural stem cells confer unique pinwheel architecture to the ventricular surface in neurogenic regions of the adult brain. Cell Stem Cell 3, 265–278.

Mizrak, D., Levitin, H.M., Delgado, A.C., Crotet, V., Yuan, J., Chaker, Z., Silva-Vargas, V., Sims, P.A., and Doetsch, F. (2019). Single-Cell Analysis of Regional Differences in Adult V-SVZ Neural Stem Cell Lineages. Cell Rep 26, 394–406 e395.

Nagayama, S., Homma, R., and Imamura, F. (2014). Neuronal organization of olfactory bulb circuits. Front Neural Circuits 8, 98.

Obernier, K., and Alvarez-Buylla, A. (2019). Neural stem cells: origin, heterogeneity and regulation in the adult mammalian brain. Development 146.

Obernier, K., Cebrian-Silla, A., Thomson, M., Parraguez, J.I., Anderson, R., Guinto, C., Rodas Rodriguez, J., Garcia-Verdugo, J.M., and Alvarez-Buylla, A. (2018). Adult Neurogenesis Is Sustained by Symmetric Self-Renewal and Differentiation. Cell Stem Cell 22, 221–234 e228.

Ortega, F., Gascon, S., Masserdotti, G., Deshpande, A., Simon, C., Fischer, J., Dimou, L., Chichung Lie, D., Schroeder, T., and Berninger, B. (2013). Oligodendrogliogenic and neurogenic adult subependymal zone neural stem cells constitute distinct lineages and exhibit differential responsiveness to Wnt signalling. Nat Cell Biol 15, 602–613.

Pentinmikko, N., Iqbal, S., Mana, M., Andersson, S., Cognetta, A.B., 3rd, Suciu, R.M., Roper, J., Luopajarvi, K., Markelin, E., Gopalakrishnan, S., et al. (2019). Notum produced by Paneth cells attenuates regeneration of aged intestinal epithelium. Nature.

Qu, Q., Sun, G., Li, W., Yang, S., Ye, P., Zhao, C., Yu, R.T., Gage, F.H., Evans, R.M., and Shi, Y. (2010). Orphan nuclear receptor TLX activates Wnt/beta-catenin signalling to stimulate neural stem cell proliferation and self-renewal. Nat Cell Biol 12, 31–40; sup pp 31-39.

R. Jones, A., Forero-Vargas, M., Withers, S.P., Smith, R.S., Traas, J., Dewitte, W., and Murray, J.A.H. (2017). Cell-size dependent progression of the cell cycle creates homeostasis and flexibility of plant cell size. Nat Commun 8, 15060.

Schindelin, J., Arganda-Carreras, I., Frise, E., Kaynig, V., Longair, M., Pietzsch, T., Preibisch, S., Rueden, C., Saalfeld, S., Schmid, B., et al. (2012). Fiji: an open-source platform for biological-image analysis. Nat Methods 9, 676–682.

Shapiro, L.A., Ng, K., Zhou, Q.Y., and Ribak, C.E. (2009). Subventricular zone-derived, newly generated neurons populate several olfactory and limbic forebrain regions. Epilepsy Behav 14 Suppl 1, 74–80.

Shekhar, K., Lapan, S.W., Whitney, I.E., Tran, N.M., Macosko, E.Z., Kowalczyk, M., Adiconis, X., Levin, J.Z., Nemesh, J., Goldman, M., et al. (2016). Comprehensive Classification of Retinal Bipolar Neurons by Single-Cell Transcriptomics. Cell 166, 1308–1323 e1330.

Stoeckius, M., Hafemeister, C., Stephenson, W., Houck-Loomis, B., Chattopadhyay, P.K., Swerdlow, H., Satija, R., and Smibert, P. (2017). Simultaneous epitope and transcriptome measurement in single cells. Nat Methods 14, 865–868.

Subramanian, A., Tamayo, P., Mootha, V.K., Mukherjee, S., Ebert, B.L., Gillette, M.A., Paulovich, A., Pomeroy, S.L., Golub, T.R., Lander, E.S., et al. (2005). Gene set enrichment analysis: a knowledge-based approach for interpreting genome-wide expression profiles. Proc Natl Acad Sci U S A 102, 15545–15550.

Suciu, R.M., Cognetta, A.B., 3rd, Potter, Z.E., and Cravatt, B.F. (2018). Selective Irreversible Inhibitors of the Wnt-Deacylating Enzyme NOTUM Developed by Activity-Based Protein Profiling. ACS Med Chem Lett 9, 563–568.

Tepe, B., Hill, M.C., Pekarek, B.T., Hunt, P.J., Martin, T.J., Martin, J.F., and Arenkiel, B.R. (2018). Single-Cell RNA-Seq of Mouse Olfactory Bulb Reveals Cellular Heterogeneity and Activity-Dependent Molecular Census of Adult-Born Neurons. Cell Rep 25, 2689–2703 e2683.

Van der Maaten, L., and Hinton, G. (2008). Visualizing Data using t-SNE. Journal of Machine Learning Research 9, 2579–2605.

Weinreb, C., Wolock, S., and Klein, A. (2017). SPRING: a kinetic interface for visualizing high dimensional single-cell expression data. Bioinformatics.

Wen, Y., Zhang, Z., Li, Z., Liu, G., Tao, G., Song, X., Xu, Z., Shang, Z., Guo, T., Su, Z., et al. (2019). The PROK2/PROKR2 signaling pathway is required for the migration of most olfactory bulb interneurons. J Comp Neurol.

Wojcinski, A., Lawton, A.K., Bayin, N.S., Lao, Z., Stephen, D.N., and Joyner, A.L. (2017). Cerebellar granule cell replenishment postinjury by adaptive reprogramming of Nestin(+) progenitors. Nat Neurosci 20, 1361–1370.

Yuan, J., Sheng, J., and Sims, P.A. (2018). SCOPE-Seq: a scalable technology for linking live cell imaging and single-cell RNA sequencing. Genome Biol 19, 227.

Yuan, J., and Sims, P.A. (2016). An Automated Microwell Platform for Large-Scale Single Cell RNA-Seq. Sci Rep 6, 33883.

Zhang, X., Cheong, S.M., Amado, N.G., Reis, A.H., MacDonald, B.T., Zebisch, M., Jones, E.Y., Abreu, J.G., and He, X. (2015). Notum is required for neural and head induction via Wnt deacylation, oxidation, and inactivation. Dev Cell 32, 719–730.

Zhang, Y., Chen, K., Sloan, S.A., Bennett, M.L., Scholze, A.R., O’Keeffe, S., Phatnani, H.P., Guarnieri, P., Caneda, C., Ruderisch, N., et al. (2014). An RNA-sequencing transcriptome and splicing database of glia, neurons, and vascular cells of the cerebral cortex. J Neurosci 34, 11929–11947.

Zheng, G.X., Lau, B.T., Schnall-Levin, M., Jarosz, M., Bell, J.M., Hindson, C.M., Kyriazopoulou-Panagiotopoulou, S., Masquelier, D.A., Merrill, L., Terry, J.M., et al. (2016). Haplotyping germline and cancer genomes with high-throughput linked-read sequencing. Nat Biotechnol 34, 303–311.

Zweifel, S., Marcy, G., Lo Guidice, Q., Li, D., Heinrich, C., Azim, K., and Raineteau, O. (2018). HOPX Defines Heterogeneity of Postnatal Subventricular Zone Neural Stem Cells. Stem Cell Reports 11, 770–783.

Zywitza, V., Misios, A., Bunatyan, L., Willnow, T.E., and Rajewsky, N. (2018). Single-Cell Transcriptomics Characterizes Cell Types in the Subventricular Zone and Uncovers Molecular Defects Impairing Adult Neurogenesis. Cell Rep 25, 2457–2469 e2458.

